# Gluconeogenesis and glycogen metabolism in the epidermis and endoderm of *Xenopus tropicalis* embryos and larvae

**DOI:** 10.64898/2026.05.08.723674

**Authors:** Motoharu Aoki, Ayaka Tsuchida, Kei Tamura, Otto Baba, Kazutoshi Yoshitake, Fumiya Furukawa

## Abstract

In many oviparous animals, egg yolk is the sole source of nutrition until feeding begins, and carbohydrates are present in only small amounts in the yolk. Glucose plays an important role in the developmental processes of various animals. In addition, gluconeogenesis has been reported to occur in the yolk syncytial layer (YSL) of cartilaginous fish and teleosts. In contrast, the role of gluconeogenesis in tetrapods remains unclear. In this study, we used *Xenopus tropicalis*, an anuran amphibian, which lacks YSL, and therefore provide an opportunity to examine the evolutionary conservation of gluconeogenic mechanisms among vertebrates. In *X. tropicalis*, liquid chromatography/mass spectrometry revealed that glucose levels increased before liver formation. Subsequent tracer experiments using ^13^C-labeled metabolic substrates detected gluconeogenesis activity from glycerol and lactate. Expression analyses showed that gluconeogenic genes are expressed in the epidermis and endoderm. Consistently, generation 0 (G0) knockout of *fbp1*, a key gluconeogenic gene, resulted in a significant reduction in glucose levels, affecting brain development. These findings first demonstrate that gluconeogenesis supports development of *X. tropicalis*. To the best of our knowledge, gluconeogenesis in developing epidermis has not been reported, highlighting previously unrecognized diversity in tissue-specific metabolism during vertebrate development. Comparative analyses across species will provide further insights into the evolution and functional significance of embryonic gluconeogenesis and nutrient metabolism.

## Introduction

In oviparous animals, egg yolk is the only source of nutrition until feeding begins. However, how embryos of such animals absorb and metabolize egg yolk remains largely unknown. The yolks of many oviparous animals contain a large amount of protein and lipids, which have been the focus of much research (Territo and Smits, 1998; Vastag, 2011). On the other hand, carbohydrates and sugars, such as glucose, have not been considered important because they are present in the yolk to a very small amount (Needham, 1927). However, in zebrafish (*Danio rerio*), knocking down a glucose transporter (Glut1) that facilitates glucose uptake into cells resulted in abnormal formation of the brain and hematopoietic stem cells in embryos (Jensen et al., 2006; Harris et al., 2013). Furthermore, in chick embryos, an oviparous tetrapod, inhibition of glycolysis using glucose analog caused arrest of body axis elongation and defects in pattern formation (Oginuma et al., 2017). These findings suggest that glucose plays important roles in the development of the zebrafish and chick embryos.

Glucose is produced by the breakdown of glycogen and can also be synthesized from amino acids, glycerol, and lactate through a metabolic pathway known as gluconeogenesis (Agius, 2015; Holeček., 2023). In vertebrates, gluconeogenesis is known to occur primarily in the liver and kidneys, where the synthesized glucose is released into the bloodstream and distributed throughout the body as an energy source via glycolysis (Sahoo et al., 2023). In addition, glucose is diverted into the pentose phosphate pathway (PPP) via glucose-6-phosphate (G6P), thereby providing precursors for nucleotide and lipid biosynthesis that support cell proliferation and differentiation (Stincone et al., 2014; DeBerardinis et al., 2007). Furthermore, glucose flux through the hexosamine biosynthetic pathway (HBP) contributes to glycosylation and the maturation of secretory proteins (Chaveroux et al., 2016; Denzel and Antebi, 2015). Given these multifaceted roles of glucose, it is reasonable that it takes important roles in developmental processes of animals.

Although glucose was found necessary during development of some animals, how their embryos obtained glucose, which is trace in egg yolks, has remained largely unclear. Recently, gluconeogenesis during embryonic development has been reported in the yolk syncytial layer (YSL), a unique extraembryonic tissue, of teleost fishes such as zebrafish and the marine grass puffer (*Takifugu niphobles*), as well as in YSL-like tissue of the cartilaginous fish, cloudy catshark (*Scyliorhinus torazame*) (Furukawa et al., 2024; Kodama et al., 2024; Shimizu et al., 2024). In these animals, gluconeogenic activities shifted to livers after development of this tissue, and such temporal extrahepatic gluconeogenesis was proposed to be unique to early developmental stages. Furthermore, endodermal tissues of the basal actinopterygian fish, sterlet (*Acipenser ruthenus*), also expressed gluconeogenic genes, suggesting that this activity is not restricted to YSL or YSL-like tissues, but is an evolutionarily conserved metabolic feature that a wide range of vertebrate lineages may exhibit in different tissues (Shimizu et al., 2025). Meanwhile, similar activities have not been reported in tetrapod lineages, leaving open the question of whether developmental extrahepatic gluconeogenesis is limited to sharks and actinopterygian animals or conserved in the wider range of vertebrate species.

In the present study, we focused on *Xenopus tropicalis* (*X. tropicalis*), or western clawed frog, an anuran amphibian species. This species is closely related to *Xenopus laevis*, or African clawed frog, a classical model organism in developmental biology (Harland and Gerhart, 1997; Nieuwkoop and Faber, 1994; Zahn et al., 2022; Harland and Grainger, 2011), and has been widely used as an experimental system suitable for genetic and genomic analysis (Hellsten et al., 2010; Grainger, 2012). Furthermore, this species exhibits holoblastic cleavage, and the yolk is stored intracellularly as yolk platelets within endodermal cells, being subsequently degraded during development (Shah et al., 2024). These features contrast with those in the sterlet, in which yolk-rich cells are located extraembryonically and function as extraembryonic tissue, as observed in teleost and cartilaginous fishes (Shah et al., 2024). Therefore, this species serves as a useful model for comparative studies against actinopterygian fishes and sharks.

In this study, we investigated gluconeogenesis and glycogen metabolism during development in *X. tropicalis*. First, we analyzed developmental changes in glucose, glycogen, and other intermediary metabolites by liquid chromatography (LC)-mass spectrometry (MS). Next, we performed isotope tracing using ^13^C-labeled metabolic substrates to directly detect gluconeogenic activities. We further examined spatiotemporal expression profiles of genes involved in gluconeogenesis and glycogen metabolism. Finally, we generated *fbp1* generation 0 (G0) knockout (KO) larvae and assessed its effect on gluconeogenic activity and brain development. The series of experiments provides not only a previously unrecognized metabolic feature of *X. tropicalis* larvae, but also a new insight into the evolutionary diversification of gluconeogenic mechanisms during vertebrate development.

## Results

### Gluconeogenic metabolites increased during hatching and liver formation

Metabolomic profiling of developing *X. tropicalis* using LC-MS revealed that the levels of many metabolites increased as development progressed (Fig. 1). Notably, glucose levels increased from 24 to 48-hours post fertilization (hpf), a period immediately after hatching (Showell and Conlon, 2013). Measurement of glycogen content revealed a marked increase between 48 and 72 hpf, a period corresponding to the liver formation stage (Supplementary Fig. S1; Zahn et al., 2022). In addition, glycerol-3-phosphate (G3P), a phosphorylated form of glycerol and a key substrate for gluconeogenesis, showed an increasing trend from 24 hpf onward. Lactate and alanine levels also increased during this period. Furthermore, tricarboxylic acid (TCA) cycle intermediates, including malate, fumarate and citrate, exhibited accumulating patterns similar to that of glucose. Since the embryos and larvae were cultured without feeding, these results suggest that gluconeogenesis is actively engaged during *X. tropicalis* development, particularly from the hatching stage (24 hpf) to the liver formation stage (72 hpf) (Supplementary Fig. S1; Zahn et al., 2022).

**Fig. 1.**
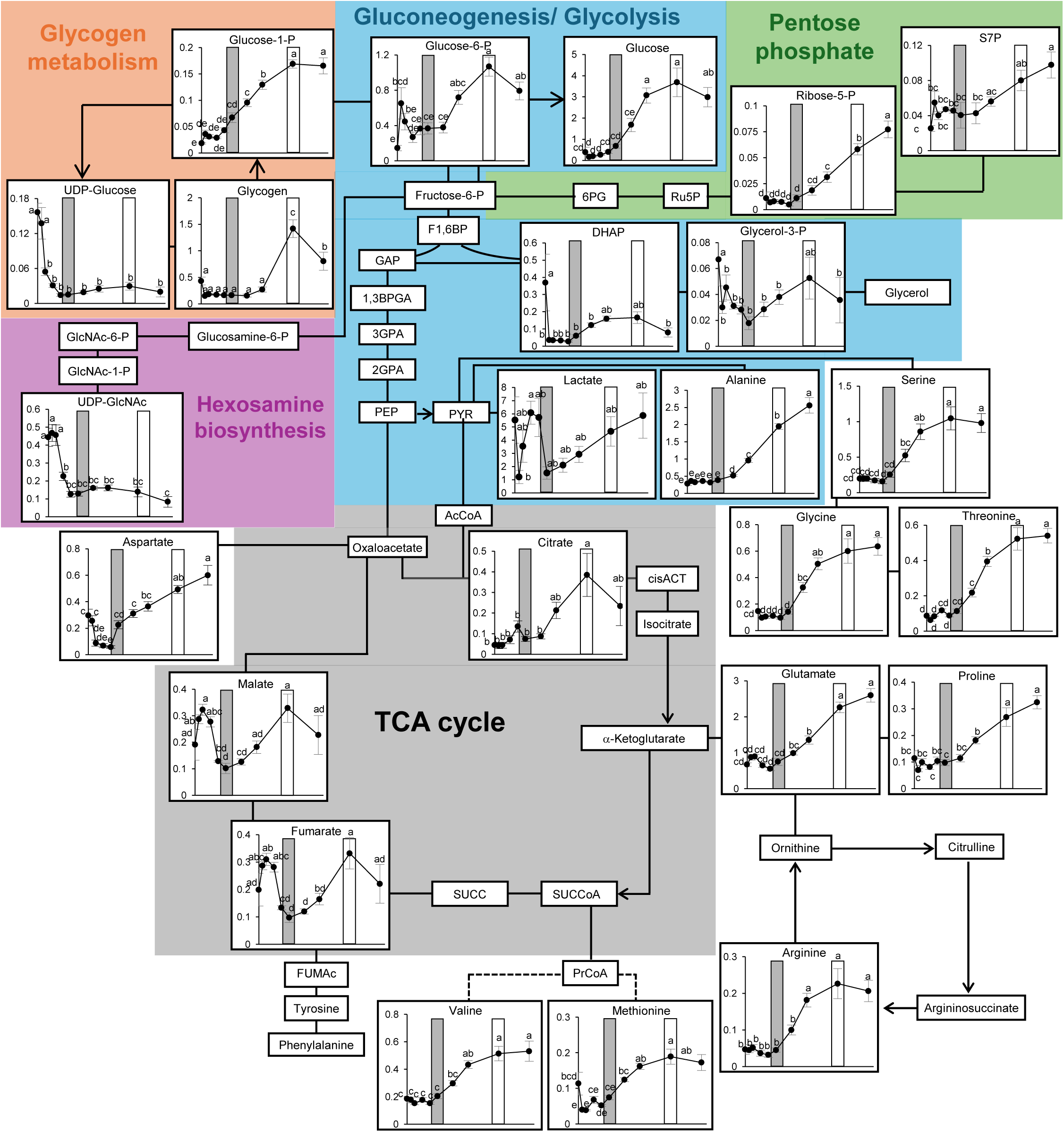
Changes in metabolite levels during development of *X. tropicalis*. The horizontal and vertical axes represent hours post-fertilization (hpf: 0, 3, 6, 12, 18, 24, 36, 48, 72 and 96) and metabolite contents (nmol/individual), respectively. In each graph, hatching (gray shading, 24 hpf) and liver-formation (white box, 72 hpf) periods are indicated. Data are presented as mean ±SE (*n*=5-7). Different letters indicate significant differences among time points (*p*<0.05), as determined by one-way analysis of variance (ANOVA) and following Tukey’s multiple comparison test. Glucose-1-P, Glucose-1-phosphate; Glucose-6-P, Glucose-6-phosphate; 6PG, 6-phosphogluconicacid; Ru5P, Ribulose-5-phosphate; Ribose-5-P, Ribose-5-phosphate; S7P, Sedoheptulose-7-phosphate; Fructose-6-P, Fructose-6-phosphate; Glucosamine-6-P, Glucosamine-6-phosphate; GlcNAc-6-P, N-acetylglucosamine-6-phosphate; GlcNAc-1-P, N-acetylglucosamine-1-phosphate; UDP-GlcNAc, UDP-N-acetylglucosamine; F1,6BP, Fructose-1,6-phosphate; DHAP, Dihydroxyacetone Phosphate; Glycerol-3-P, Glycerol-3-phosphate; GAP, Glyceraldehyde-3-phosphate; 1,3BPGA, 1,3-Bisphosphoglycerate; 3GPA, 3-phosphoglycerate; 2GPA, 2-phosphoglycerate; PEP, Phosphoenolpyruvate; PYR, Pyruvate; AcCoA, Acetyl-coenzyme; cisACT, cis-aconitate; SUCC, Succinate; SUCCoA, Succinyl-coenzyme; PrCoA, Pionyl-coenzyme; FUMAc, Fumarylacetoacetate.

### Larval gluconeogenesis occurs from lactate and glycerol as substrates

To determine whether gluconeogenesis actually occurs during development, we performed metabolite tracing analysis using ^13^C-labeled alanine, lactate, glutamine, glutamate and glycerol (Fig. 2). 24-hpf *X. tropicalis* larvae were cut into thirds and cultured for 6 h in media supplemented with ^13^C-labeled substrates (Fig. 2 A). If ^13^C-labeled substrates are metabolized through the gluconeogenesis pathway and converted into glucose, the resulting glucose is detected as a mass +3 (M+3) isotopologue, containing three ^13^C atoms (Fig. 2 B). As a result of this experiment, glucose M+3 significantly increased when supplemented with lactate-^13^C_3_ or glycerol-^13^C_3_ as substrates (Fig. 2 C, supplementary Fig. S2). In addition, a gluconeogenic intermediate G6P M+3 was also detected under all substrate conditions, with particularly high labeling observed in larvae treated with glycerol-^13^C_3_. These results demonstrate gluconeogenic activity in the *X. tropicalis* larvae. M+3 isotopologues of glucose-1-phosphate (G1P) and UDP-glucose, the intermediates involved in glycogen metabolism, also increased under conditions containing ^13^C-labeled substrates (Fig. 2 B and C, supplementary Fig. S2). This result suggests that glycogen synthesis is concurrently activated during periods of enhanced gluconeogenesis, while the contribution of glycogen breakdown to glucose supply is likely to be minor. Meanwhile, M+3 isotopologues of sedoheptulose-7-phosphate (S7P), an intermediate of the PPP, and UDP-*N*-acetylglucosamine (UDP-GlcNAc), a key metabolite in the HBP, also increased under all ^13^C substrate conditions (Fig. 2 B and C, supplementary Fig. S2). These findings indicate that metabolites produced via gluconeogenesis are not only used for energy metabolism but also flow into biosynthetic pathways, including the pentose phosphate and hexosamine biosynthesis pathways. TCA cycle is a key metabolic pathway that incorporates the carbon derived from glucogenic amino acids (e.g. glutamine, glutamate, and alanine) as well as lactate toward gluconeogenesis (Anderson et al., 2018; Hui et al., 2017). This cycle maintains its carbon pool by balancing the influx and efflux of intermediates through anaplerotic and cataplerotic reactions, respectively (Inigo et al., 2021). We expected that our metabolic tracing with ^13^C-labeled substrates generates isotopologues with varied numbers of ^13^C. To capture these dynamics, we analyzed labeled metabolites ranging from M+2 to M+6 (Fig. 2 B). Under lactate-^13^C_3_ supplementation, high labeling of the M+2 and M+3 isotopologues was observed for citrate, α-ketoglutarate, succinate, and malate (Fig. 2 D, Supplementary Fig. S3). Also, higher labeling of the M+4 and M+5 isotopologues was detected under glutamate-^13^C_5_ supplementation (Fig. 2 D, Supplementary Fig. S3). In contrast, glycerol-^13^C_3_ did not produce any labeled TCA cycle isotopologues assessed in the present study. These results suggest that lactate and glutamine/glutamate, but not glycerol, serve as major carbon sources fueling the TCA cycle in *X. tropicalis* larvae. In humans, mice, and zebrafish, lactate, glutamine and/or glutamate act as major carbon donors to the TCA cycle (DeBerardinis et al., 2007; Hui et al., 2017; Furukawa et al., 2024), suggesting that such metabolic framework is common in many vertebrate animals including *X. tropicalis*.

**Fig. 2.**
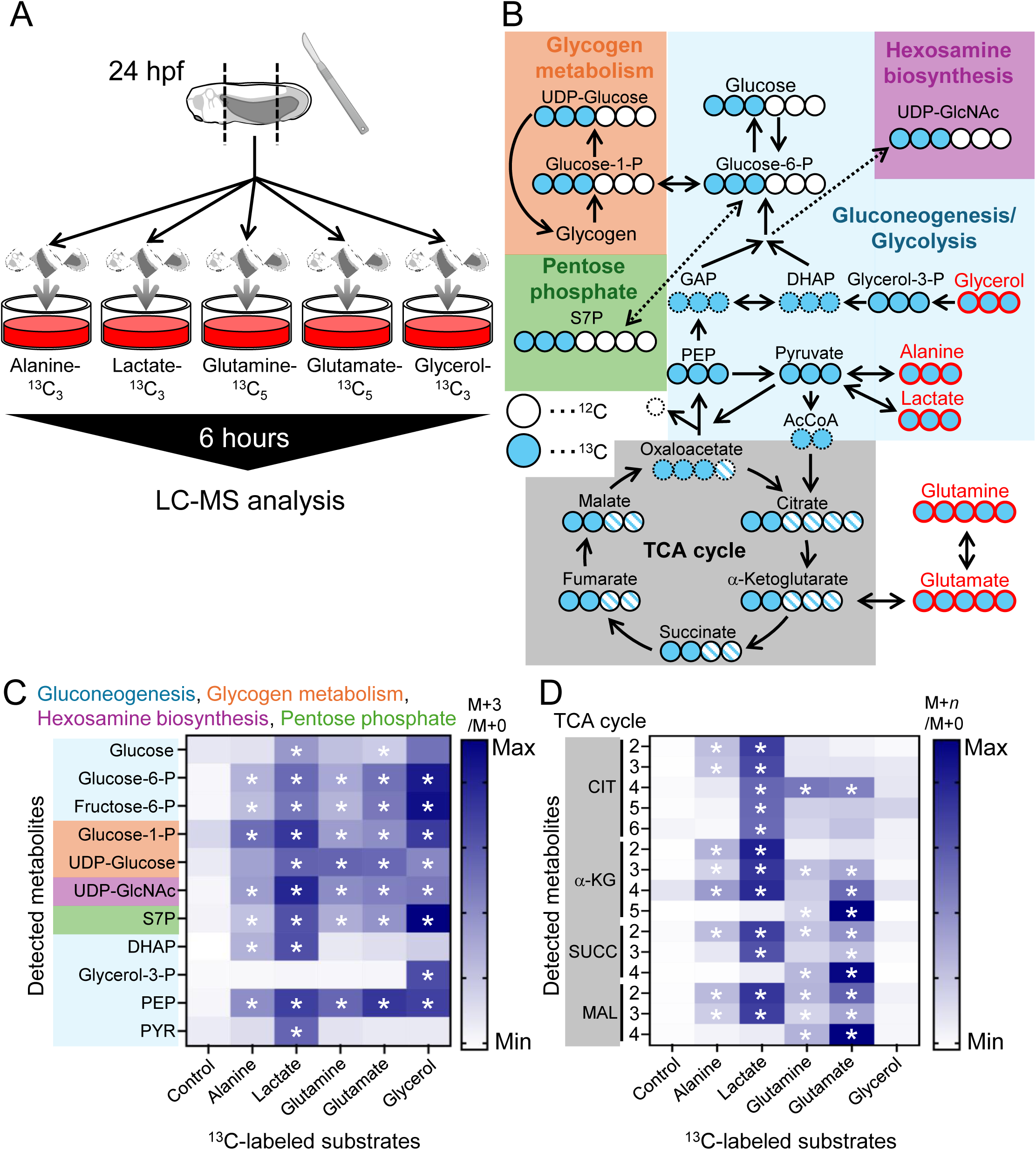
LC-MS-based isotope tracing reveals gluconeogenesis activity in *X. tropicalis* larvae. (A) Schematic overview of the gluconeogenic pathway with expected flow and incorporation of ^13^C into gluconeogenic intermediate metabolites. Open circles indicate carbon atoms (^12^C) in metabolites, and those filled with light blue indicate ^13^C. Hatched circles indicate either ^12^C or ^13^C, and dashed circles indicate metabolites not assessed. (B) 24 hpf larvae were incubated with substrates-^13^C for 6 h, followed by metabolite extraction and analysis using LC-MS. (C) A heatmap showing the ratio of mass (M) +3 to M+0 isotopologues of metabolites involved in gluconeogenesis/glycolysis, glycogen metabolism, the pentose phosphate pathway, and the hexosamine biosynthesis pathway. The horizontal axis indicates the supplied ^13^C-labeled substrates (alanine, lactate, glutamine, glutamate, and glycerol), and the vertical axis indicates the detected M+3 metabolites. (D) A heatmap showing the ratio of M+*n* to M+0 isotopologues of metabolites associated with the tricarboxylic acid (TCA) cycle. The horizontal axis indicates the supplied ^13^C-labeled substrates, and the vertical axis indicates the detected M+*n* isotopologues. *n* denotes the number of additional mass. In each block, darker blue indicates a higher proportion of the indicated isotopologues. Glucose-6-P, Glucose-6-phosphate; Fructose-6-P, Fructose-6-phosphate; Glucose-1-P, Glucose-1-phosphate; UDP-GlcNAc, UDP-N-acetylglucosamine; S7P, Sedoheptulose-7-phosphate; GAP, Glyceraldehyde-3-phosphate; DHAP, Dihydroxyacetone Phosphate; Glycerol-3-P, Glycerol-3-phosphate; PEP, Phosphoenolpyruvate; PYR. Pyruvate; AcCoA, Acetyl-coenzyme A; CIT, Citrate; α-KG, α-Ketoglutarate; SUCC, Succinate; MAL, Malate.

### Expression of gluconeogenic genes increases at hatching period

By quantitative PCR (qPCR) analysis, we assessed changes in the expression of genes involved in gluconeogenesis and glycogen metabolism: glucose-6-phosphatase (G6PC), *g6pc1*/*g6pc1.2*/*g6pc1.3*; glycerol-3-phosphate dehydrogenase (GPD), *gpd1/gpd1l/gpd2*, fructose-1,6-phosphatase (FBP), *fbp1*; lactate dehydrogenase (LDH), *ldha*/*ldhb*; phosphoenolpyruvate carboxykinase (PEPCK), *pck1*/*pck2*; glycogen synthase (GYS), *gys1*/*gys2*; glycogen phosphorylase (PYG), *pygb*/*pygm*/*pygl*. Expression levels of *g6pc1*/*g6pc1.2*/*g6pc1.3* and *fbp1* increased markedly between 18 and 24 hpf (Fig. 3). In addition, the expression of *ldha*, which is involved in the interconversion between lactate and pyruvate, as well as *gpd1*, which catalyzes the reversible redox reaction converting glycerol-3-phosphate to dihydroxyacetone phosphate, showed an increasing trend from 12 hpf onward (Fig. 3). These results indicate that the upregulation of gluconeogenic gene expression precedes the increase in glucose content during development (Fig. 1). Regarding glycogen metabolism, the expression levels of the ubiquitous-type isoforms, *gys1* and *pygm* (Marr et al., 2022), showed increasing trends at 24 hpf (Fig. 3), preceding the increase in glycogen content (Fig. 1). In contrast, the expression levels of *gys2* and *pygl*, the liver-type isoforms (Agius, 2015), increased toward 72 hpf (Fig. 3), in accordance with the timing of liver formation stages (Supplementary Fig. S1; Zahn et al., 2022).

**Fig. 3.**
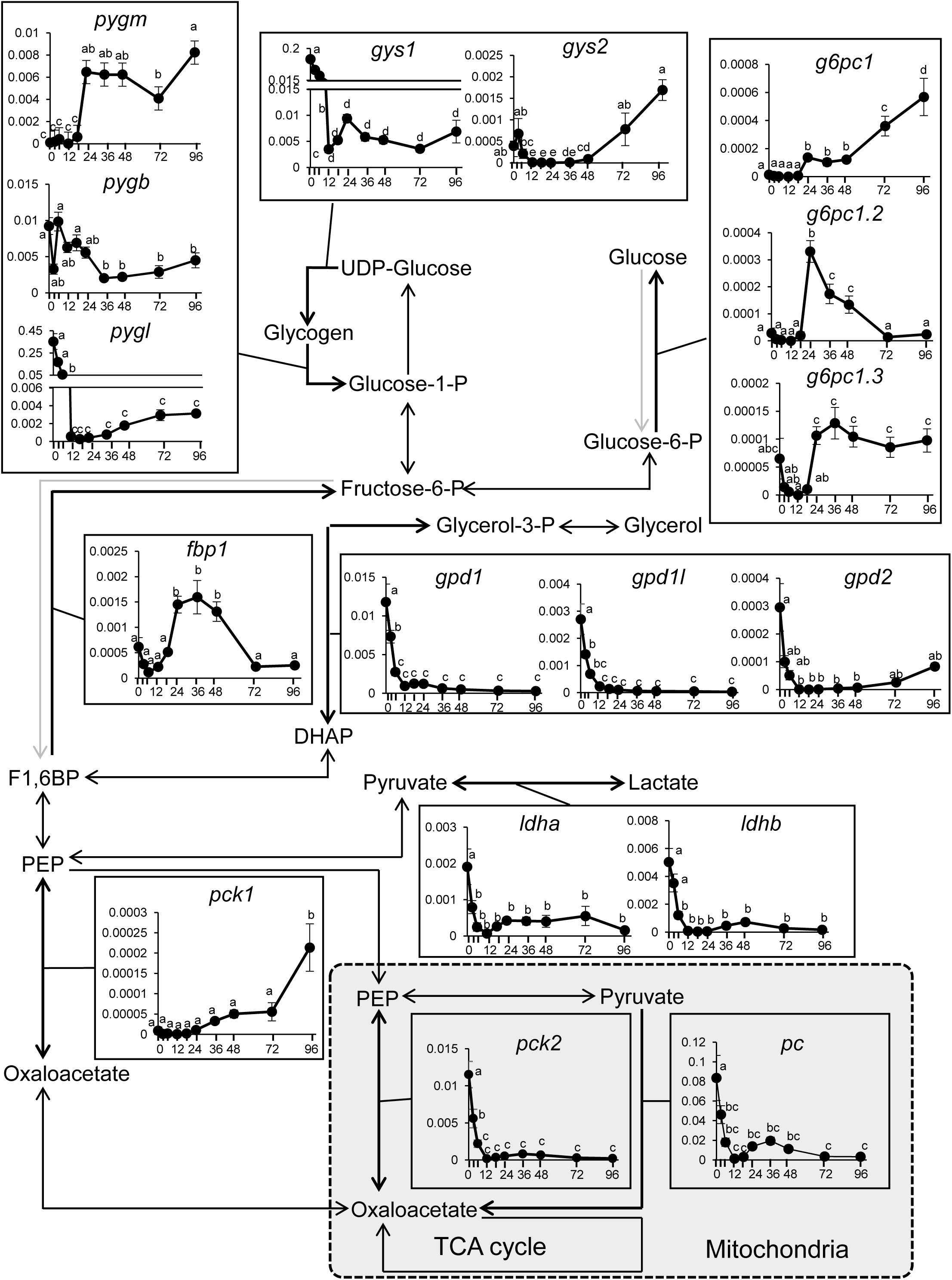
Developmental changes in the expression of gluconeogenic and glycogen metabolism genes in *X. tropicalis*. The horizontal axes represent hours post fertilization (hpf: 0, 3, 6, 12, 18, 24, 36, 48, 72 and 96), and the vertical axis represents target mRNA copy number normalized to *eef1a* copy number. Data are shown as mean ±SE (N=5-7). Time points labeled with different letters from each other indicate statistically significant difference at *p*< 0.05 determined by one-way analysis of variance (ANOVA) and following Tukey’s multiple comparison test.

### Gluconeogenic genes are expressed in the epidermis and endoderm during development

In *X. tropicalis*, and liver formation occurs at around 72 hpf (Supplementary Fig. S1; Zahn et al., 2022). Therefore, gluconeogenesis detected in the present study was expected to take place in extrahepatic tissues. We performed *in situ* hybridization analyses for gluconeogenic and glycogen metabolic genes that were examined by qPCR, using larvae at 24, 48 and 96 hpf (Fig. 4, Supplementary Figs. S1, S4-S10). At 24 hpf, the expression signals were not prominent, but those of *fbp1* and *pck1* were detected in the endoderm (Supplementary Figs. S4, S7). In the epidermis, *fbp1*, *gpd1*, *gpd1l*, *pck1*, *pck2*, and *ldhb* were also expressed (Supplementary Figs. S4, S7). At 48 hpf, the expression of most genes involved in gluconeogenesis were clearly detected: *g6pc1*, *g6pc1.2*, *g6pc1.3*, *fbp1*, *gpd1*, *pc.2*, *pck2*, and *ldhb* were expressed in endoderm (Fig. 4, Supplementary Figs. S5, S8). In addition, *g6pc1*, *g6pc1.2*, *fbp1*, *gpd1*, *pc.2*, *pck1*, *pck2*, and *ldhb* were also detected in the epidermis (Fig. 4, Supplementary Figs. S5, S8). For glycogen-metabolizing genes, *gys1*, *pygb* and *pygm*, were detected in the muscle and brain (Fig. 4, Supplementary Fig. S5). At 96 hpf, where many organs were developed, most genes assessed were expressed in the liver, intestine, and/or epidermis (Supplementary Figs. S6, S9). Additionally, *gys1*, *pygm*, and *pygb* were expressed in the muscle and/or brain, whereas *gys2* and *pygl* were expressed in liver (Fig. 4, Supplementary Fig. S6). These results suggest that gluconeogenesis occurs in endodermal and epidermal tissues of larval *X. tropicalis*. Moreover, the spatiotemporal expression patterns of *gpd1*/*gpd1l*/*gpd2* and *ldhb*, together with the result of ^13^C tracing experiment, strongly support the conclusion that lactate and glycerol are major substrates for gluconeogenesis in these tissues. In addition, glycogen synthesis appears to occur mainly in the muscle, liver, and brain, which is supported by immunofluorescence signals of glycogen in these tissues (Supplementary Fig. S10).

**Fig. 4.**
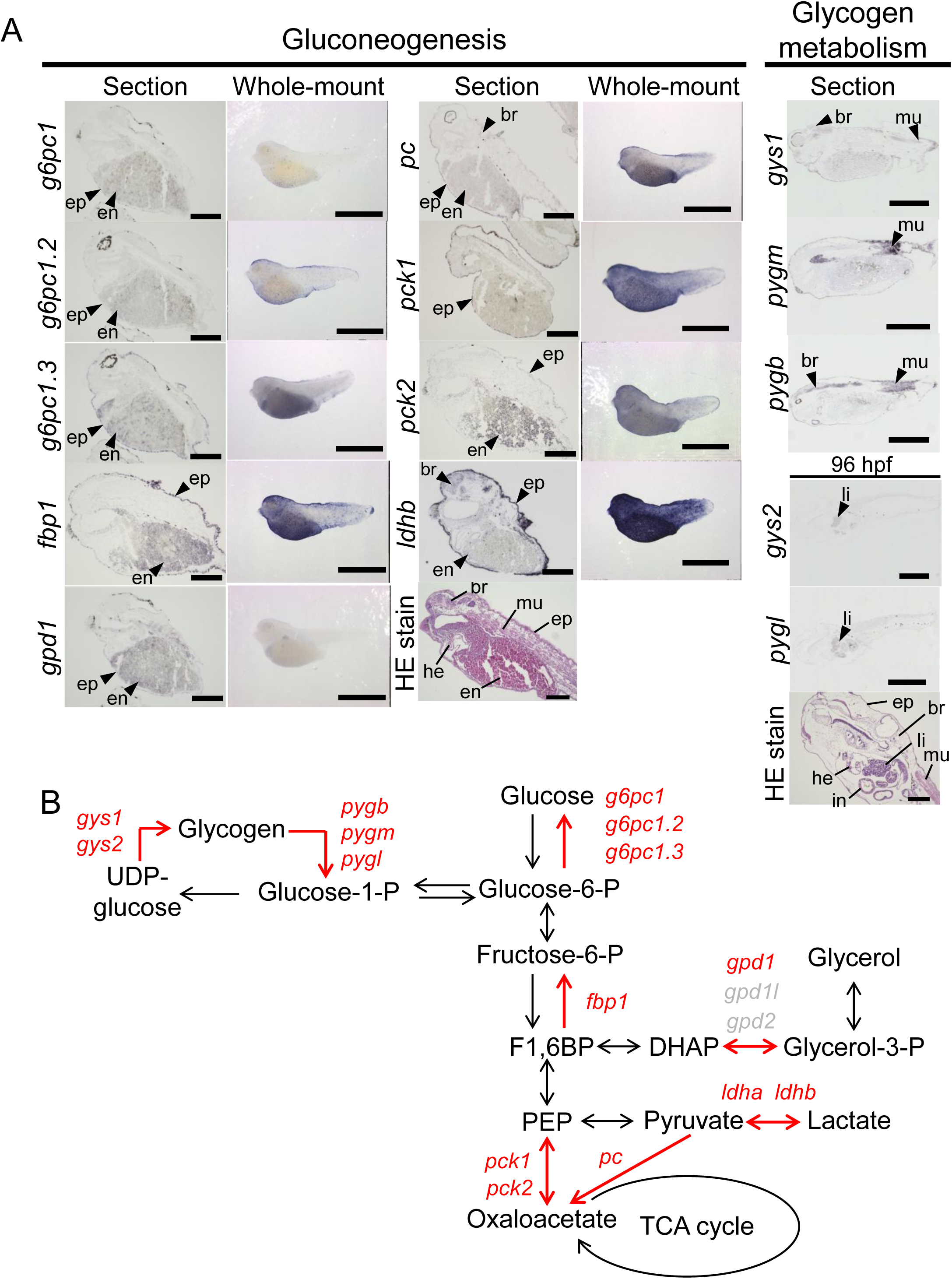
Expression of gluconeogenesis in epidermis and endoderm at 48 and 96 hpf. (A) Spatial expression patterns of gluconeogenic genes (*fbp1*, *gpd1*, *gpd1l*, *pck1, pck2*, *ldha* and *ldhb)* and glycogen metabolism genes (*gys1*, *pygb*, and *pygm*) at 48 hpf, and *gys2* and *pygl* at 96 hpf, analyzed by *in situ* hybridization. Positive signals are indicated by black arrowheads and tissue labels. Hematoxylin and eosin (HE) staining shows the general morphology of 48 and 96 hpf. Tissues are labeled as follows: br, brain; en, endoderm; ep, epidermis; he, heart; in, intestine; mu, muscle. Scale bars represent 250 μm (*in situ* hybridization on sections, gluconeogenic genes), 500 μm (*in situ* hybridization for glycogen metabolism genes), 1 mm (whole-mount *in situ* hybridization), and 200 μm (HE staining). (B) Expression of gluconeogenic and glycogen metabolism genes were mapped onto the gluconeogenesis pathway. Red letters denote the gene transcripts detected by *in situ* hybridization analysis and red arrowheads indicate the pathways that the detected genes mediate.

### Effect of *fbp1* G0 Knockout on glucose production

To examine the contribution of gluconeogenesis during development, G0 KO larvae targeting *fbp1* were generated using the CRISPR/Cas9 system, and the effects on glucose production and brain development were analyzed (Fig. 5). First, we quantified the mutation frequency (indel efficiency) at one of the *fbp1* target sites using nanopore amplicon sequencing (Fig. 5 A, Exon 1). The average editing efficiency in the G0 KO group was 67.7 ± 9.9% (*n* = 7; Fig. 5 B). Although notable individual variability was observed within the G0 KO group, ranging from 25.3% to 96.4% (Fig. 5 B; Supplementary Fig. S11, S12). In contrast, the SC group showed no detectable on-target editing, and all reads were classified as WT (data not shown). To further characterize variations of mutation patterns at the individual level, we analyzed insertion/deletion (indel) patterns and their allelic frequencies in G0 KO larvae (Fig. 5 C, Supplementary Fig. S11). As a result, limited numbers of indel events accounted for a substantial proportion of edited reads; in all individuals, the top four most frequent alleles alone accounted at least 70% of all indel mutations. Sequence alignment confirmed that these mutations were concentrated around the predicted Cas9 cleavage site, consistent with non-homologous end joining (NHEJ)-mediated repair (Supplementary Fig. S12). These results indicate that our G0 KO trials successfully reduced the intact *fbp1* loci.

**Fig. 5.**
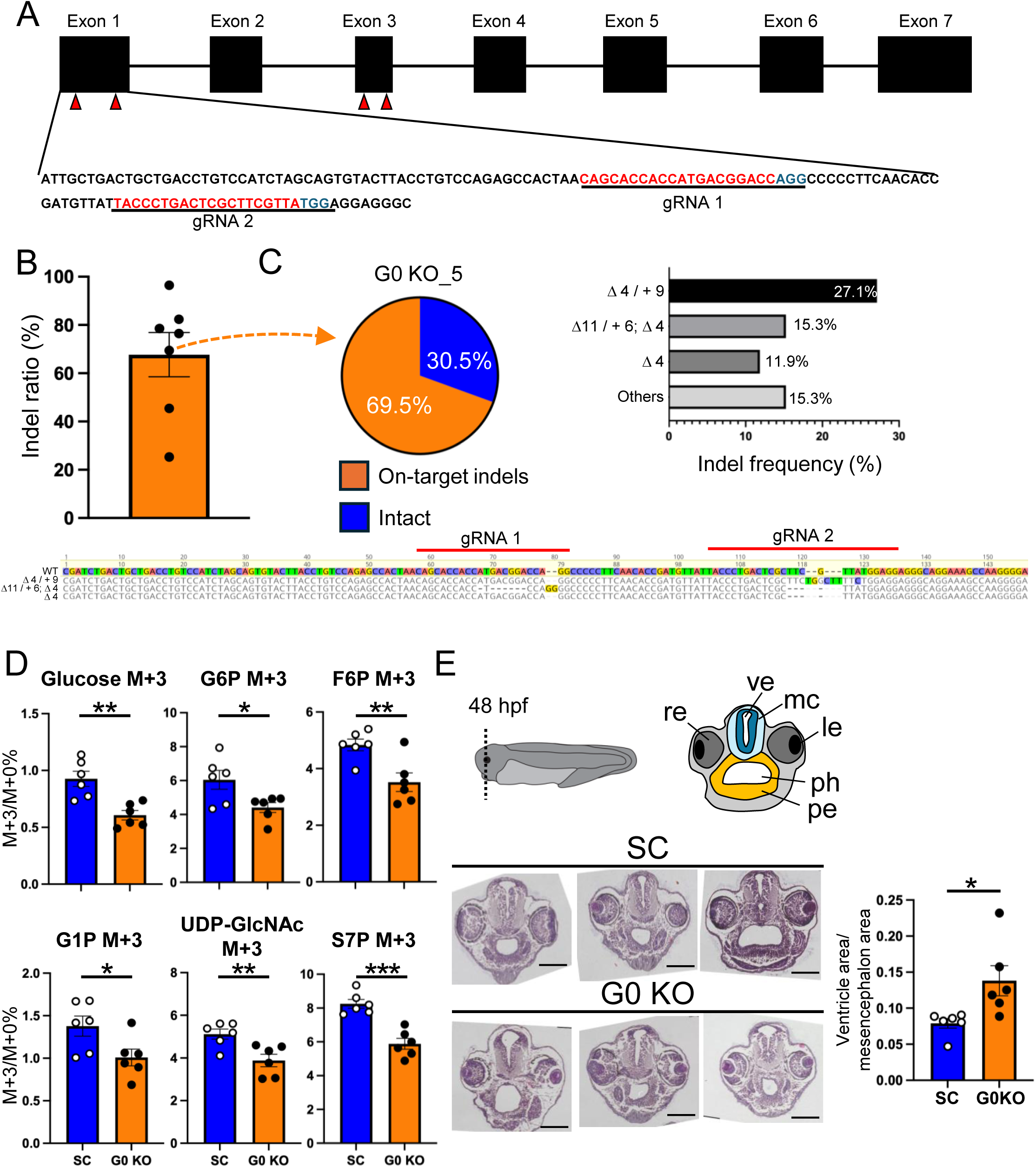
Analysis of *fbp1* in gluconeogenesis using CRISPR/Cas9 G0 KO. **(A)** Schematic diagram of the target site. The structure of the *fbp1* gene and the gRNA target sites are shown by red arrows and letters, and the PAM sequence is indicated in blue letters. **(B)** Indel efficiency KO groups. Total indel efficiency in G0 KO group (KO, n = 7) is shown. Each dot represents an individual larva, and bars indicate mean ±SE. **(C)** Allelic composition in a representative G0 KO larva (G0 KO_5). The pie chart displays the proportions of on-target indels and intact sequences. In addition, the three most frequent alleles are shown individually, with indel size and frequency (%) indicated for each allele. Deletions are indicated by Δ, while combined deletion/insertion events are presented as, for example, △ 4/ + 9. Sequences are aligned to the wild-type (WT) sequence, with deletions indicated by hyphens (–) and insertions shown in color. **(D)** Metabolomic and isotope tracing analyses in SC control and *fbp1* G0 KO larvae using LC–MS. The ratio of mass (M) +3 to M+0 isotopologues is shown for metabolites involved in gluconeogenesis/glycolysis, glycogen metabolism, the pentose phosphate pathway, and the hexosamine biosynthetic pathway. G6P, glucose-6-phosphate; G1P, glucose-1-phosphate; UDP-GlcNAc, UDP-N-acetylglucosamine; S7P, sedoheptulose-7-phosphate. The horizontal axis indicates SC and *fbp1*-targeting gRNA/Cas9-injected larvae, and the vertical axis represents the M+3/M+0 ratio of each metabolite. Each dot represents an individual replicate (5 larvae), and bars indicate mean ±SE (n = 6). Statistical comparisons between two groups were performed using Welch’s *t*-test (\**p* < 0.05, \*\**p* < 0.01, \*\*\**p* < 0.001). **(E)** Histological analysis of brain tissue by hematoxylin and eosin (HE) staining in SC and G0 KO larvae. A schematic diagram indicates the sectioning plane and corresponding brain regions. Representative HE-stained images of SC and G0 KO larvae are shown. Scale bars represent 200 μm. The graph on the right shows the ratio of ventricle area to mesencephalon area. Measurements were performed using Fiji, and bars represent mean ±SE (n = 6). Statistical comparisons between two groups were performed using Welch’s *t*-test (\**p* < 0.05). le, lens; mc, mesencephalon; pe, pharyngeal endoderm; ph, pharynx; re, retina; ve, ventricle.

Next, these *fbp1* G0 KO larvae were incubated in culture medium containing glycerol-^13^C_3_ for 6 h, followed by isotope tracing analysis. The M+3 isotopologue fractions of glucose, G6P, and F6P were decreased in the *fbp1* G0 KO group compared with the SC gRNA/Cas9-injected control group (Fig. 5 D, Supplementary Fig. S13). In addition, the M+3 fractions of S7P and UDP-GlcNAc were also decreased in the *fbp1* G0 KO group, suggesting substantial effects of reduced *fbp1* on PPP and HBP, respectively (Fig. 5 D, Supplementary Fig. S13). To evaluate the impact of reduced glucose availability on brain development, sections of 24-hpf SC and *fbp1* G0 KO larvae were stained with hematoxylin and eosin (HE) and observed under a light microscope (Fig. 5 E, Supplementary Fig. S14). As a result, the ratio of ventricle area to mesencephalon area was significantly increased in *fbp1* G0 KO larvae compared with SC larvae, suggesting that gluconeogenesis is important for proper brain development.

## Discussion

Glucose serves not only as a central substrate for energy metabolism but is also essential for critical events in early development, including cell proliferation and differentiation (Liu et al., 2025; Pantaleon et al., 2008). Indeed, in zebrafish, grass puffer, cloudy catshark, and sterlet, glucose has been reported to be synthesized via gluconeogenesis during development (Furukawa et al., 2024; Kodama et al., 2023; Shimizu et al., 2024; Shimizu et al., 2025). In this study, we aimed to understand if gluconeogenesis also occurs during development in the amphibian tetrapod *X. tropicalis* and if the underlying mechanisms are evolutionarily conserved in vertebrates including amphibians. Also, by G0 KO experiment, we assessed potential importance of gluconeogenesis in brain development of this animal.

Metabolome analysis using LC-MS revealed that glucose levels increased from 24 hpf to 48 hpf (Fig. 1), the timing well before liver formation (72 hpf). Because feeding in *X. tropicalis* begins between 72 and 96 hpf (Zahn et al., 2022), and all experiments were conducted under unfed conditions, the observed increase in glucose is most likely attributable to endogenous glucose production rather than dietary intake. In contrast, glycogen, the major storage form of glucose, remained constant until 48 hpf (Fig.1), suggesting that the early increase in glucose is unlikely to be derived from glycogen breakdown. Excluding feeding and glycogen breakdown, the glucose was most likely synthesized via gluconeogenesis.

To directly test gluconeogenic activity, we performed metabolic tracing experiments using ^13^C-labeled metabolic precursors (Fig. 2 A, B). Incorporation of ^13^C into glucose was detected following the introduction of lactate-^13^C_3_ and glycerol-^13^C_3_ (Fig. 2 C, Supplementary Fig. S2), demonstrating that *X. tropicalis* larvae actively perform gluconeogenesis using these substrates. In *X. tropicalis* eggs, lipids constitute the majority of stored nutrients (Needham, 1927), and in the closely related species *Xenopus laevis*, triacylglycerol levels decrease substantially during early embryogenesis (Rizzo et al., 1994). These observations suggest that glycerol derived from glycerolipid degradation is available as a gluconeogenic substrate in *X. tropicalis* larvae. Lactate is also abundant in unfertilized eggs and early embryos, and previous studies have shown that lactate pools contribute to carbohydrate metabolism during early development (Dworkin and Dworkin-Rastl, 1990), indicating that lactate likely serves as another major gluconeogenic substrate. In contrast, previous studies have shown that glutamate is preferentially used for gluconeogenesis in zebrafish, whereas glycerol is the dominant substrate in cloudy catshark embryos (Furukawa et al., 2024; Shimizu et al., 2024). Teleost yolks are rich in protein, while shark embryos consume a substantial fraction of yolk lipids during development (Wrisez et al., 1993; Lechenault et al., 1993). This difference suggests that preferred gluconeogenic substrates vary among taxa and reflect species-specific yolk nutrient composition.

Our results of high ^13^C-labeling rates in the glycogen synthesis pathway suggest that glycogen synthesis is promoted concurrently during periods when gluconeogenesis is highly active. Since net glycogen levels did not change during this period, the rate of synthesis and degradation of glycogen may be balanced. High ¹³C-labeling rates were also observed in intermediates of the PPP, suggesting that glucose or G6P synthesized via gluconeogenesis is also directed into the PPP. This pathway provides ribose-5-phosphate for nucleotide synthesis and to generate NADPH required for biosynthetic processes (Stincone et al., 2015; Vander Heiden et al., 2009; Patra and Hay, 2014). In addition, it has been reported that metabolic flux through this pathway promotes cell proliferation during tail regeneration in *X. tropicalis* (Patel et al., 2022). Based on these findings, during development, the PPP may be activated to utilize glucose synthesized via gluconeogenesis to support rapid cell proliferation. In addition, high labeling rates were also observed in metabolites associated with the HBP, suggesting gluconeogenesis-mediated ^13^C flow into the HBP. UDP-GlcNAc and UDP-*N*-acetyl-D-galactosamine produced through HBP serve as substrates for *O*-linked glycan, such as mucins, the major components of mucus (Bennett et al., 2012; Bansil and Turner, 2018). In *X. tropicalis*, the period between 24 and 48 hpf corresponds to the hatching stage (Showell and Conlon, 2013; Zahn et al., 2022), when larvae begin to contact the external environment. At this stage, secretory cells, including goblet and small secretory cells, are present in the epidermis (Billett and Gould, 1971; Dubaissi et al., 2014) and contribute to the formation of a physical barrier against pathogens by secreting mucins and other protective factors (Dubaissi et al., 2017). Taken together, our findings imply that glucose synthesized via gluconeogenesis is also supplied to glycan biosynthesis through HBP and play important roles in survival of larvae. Finally, high levels of ¹³C labeling were also observed in intermediates of the TCA cycle, suggesting that these substrates serve as carbon sources for mitochondrial metabolism in *X. tropicalis* larvae. This is consistent with previous reports demonstrating that lactate and glutamine/glutamate contribute substantially to the TCA cycle in proliferating cells (DeBeradinis et al., 2007; Hui et al., 2017; Furukawa et al., 2024). In contrast, alanine, glutamine, and glutamate showed low labeling efficiency for glucose M+3. This finding implies that the cells metabolizing these amino acids are distinct from those undergoing glucose production. In other words, metabolic functions may be partitioned among different cell types during development (DeBerardinis and Chandel, 2016; Bulusu et al., 2017).

Analysis of gluconeogenic gene expression revealed that the expression levels of *g6pc1*/*g6pc1.2*/*g6pc1.3*, and *fbp1* increased between 18 and 24 hpf, preceding the observed increase in glucose levels (Fig. 3). Furthermore, most gluconeogenic genes were predominantly expressed in endoderm at 24 hpf and 48 hpf (Fig. 4, Supplementary Fig. S4, S5), whereas their expression was detected in the liver and intestine at 96 hpf (Supplementary Fig. S6). These results suggest that, in *X. tropicalis*, gluconeogenesis occurs in yolk-rich endoderm first, and then in endoderm-derived organs such as the liver and intestine. These expression patterns are comparable to those observed in sterlet, where yolk sac endoderm expressed gluconeogenic genes, and the expression is maintained in the same region even after differentiation into the intestine and liver (Shimizu et al., 2025). In contrast, in zebrafish, grass puffer and cloudy catshark, temporary gluconeogenesis has been reported to occur in extraembryonic tissues such as the YSL or YSL-like structures (Furukawa et al., 2024; Kodama et al., 2024; Shimizu et al., 2024), utilizing yolk-derived nutrients. Since these extraembryonic tissues regress as the development proceeds (Kimmel et al., 1985; Trinkaus et al., 1993), the primary site of gluconeogenesis shifts to the liver. These interspecies differences are likely attributable to differences in cleavage patterns and yolk distribution. In amphibians and sterlet, holoblastic cleavage represents an ancestral developmental mode in which yolk is incorporated into endodermal tissues within the embryo or extraembryonic cells (Takeuchi et al., 2008; Shah et al., 2024). In contrast, in lineages exhibiting meroblastic cleavage, the yolk is segregated into extraembryonic tissues (Takeuchi et al., 2008). This difference in nutrient supply mechanisms may explain the divergence in gluconeogenic tissues during development.

Expressions of gluconeogenic genes were also detected in the epidermis at 24, 48 and 96 hpf. (Fig. 4, Supplementary Fig. S4-S9). Gluconeogenesis in the epidermis has been reported previously: lactate-driven gluconeogenesis occurs in the outer root sheath of human hair follicles (Figlak et al., 2021; Ruan et al., 2025). In addition, in *Caenorhabditis elegans*, epidermal PCK2-dependent gluconeogenesis provides energy required for mating behavior in mid-adult males (Goncalves et al., 2020). In contrast, gluconeogenesis in the epidermis during developmental stages has not been well characterized. The epidermis of *X. tropicalis* embryos and larvae consists of diverse cell types (Walentek, 2020), including ionocytes (Quigley et al., 2011), mucus-secreting goblet cells and small secretory cells (Billett and Gould, 1971; Dubaissi et al., 2014), as well as multiciliated cells bearing numerous cilia (Assheton, 1896). These facts imply that local gluconeogenesis in the epidermis supports the activities of these functionally diverse epidermal cells.

In mammals, glycogen is predominantly stored in skeletal muscle and liver, although glycogen synthesis can occur in other tissues (Roach et al., 2012). The liver plays a central role in blood glucose homeostasis by releasing glucose derived from glycogenolysis or gluconeogenesis into the circulation and storing excess glucose as glycogen (Agius, 2015). In the present study, glycogen synthesis and degradation were observed in somite, muscle, and brain before liver formation during early development (Figs. 3, 4, Supplementary Figs. S4-S6). These findings suggest that the primary site of glycogen metabolism shifts to the liver as hepatic development proceeds. Moreover, the increase in glycogen content following the rise in glucose levels (Fig. 1), together with the synthesis of glycogen-related intermediary metabolites from gluconeogenic substrates (Fig. 2. Supplementary Fig. S2), indicates that glucose produced during development is not only consumed but is stored as glycogen. Yolk degradation progresses during development (Jorgensen et al., 2009; Fogotto and Maxfield, 1994) and feeding is initiated between 72 and 96 hpf (Zahn et al., 2022). This transition is accompanied by a reduction in yolk-derived nutrient supply. In this study, glycogen levels decreased between 72 and 96 hpf, while metabolites of the PPP showed an increasing trend during the same period (Fig. 1). These results suggest that, when the yolk becomes scarce and exogenous food is not available, stored glycogen is mobilized and utilized as a metabolic substrate. This shift likely reflects the transition from yolk-dependent nutrient supply to exogenous energy utilization.

Genome editing using the CRISPR/Cas9 system is known to induce mosaicism, in which cells with different genotypes coexist within the same individual (Mehravar et al., 2019). In this study, *fbp1* G0 KO larvae were used; therefore, the potential impact of mosaicism must be taken into consideration. Amplicon analysis by nanopore sequencing revealed an average indel efficiency of 67.7 ± 9.9% (*n* = 7) at the *fbp1* target locus in G0 KO larvae (Figs. 5 A, B; Supplementary Fig. S11). However, there was a substantial inter-individual variation in editing efficiency, which may contribute to variations in the impact of *fbp1* G0 KO among individuals. Thus, future studies using stable KO lines will be necessary to precisely evaluate the role of *fbp1*. In the metabolic tracing experiment, the loss of *fbp1* resulted in a significant reduction in glucose production from glycerol, demonstrating reduced gluconeogenic activity (Fig. 5 D; Supplementary Fig. S13). Furthermore, the production of metabolites associated with the PPP and HBP was also decreased, suggesting that glucose synthesized via *fbp1*-dependent gluconeogenesis is supplied to these downstream metabolic pathways. A histological analysis of 48 hpf larvae revealed an expansion of the brain ventricle region in *fbp1* G0 KO larvae (Fig. 5 E, Supplementary Fig. S14). Previous studies in zebrafish have shown that knockdown of glucose transporters leads to defects in ventricle formation (Jensen et al., 2006), highlighting the importance of glucose availability in central nervous system development. In *fbp1*-G0 KO larvae, the reduction in PPP flux suggests that this may have affected cell proliferation and tissue morphogenesis during development. Indeed, PPP activity is high in the early developing brain (Hakim et al., 1980), and NADPH produced through this pathway plays a critical role in maintaining redox homeostasis in neural cells (Cao et al., 2017). Therefore, a reduced supply of such PPP-derived metabolite may impair proper neural development by decreasing cell proliferation and increasing cell death.

In conclusion, this study demonstrates that gluconeogenesis occurs during the developmental process of *X. tropicalis*, most likely in the endoderm and epidermis. In amphibians, endodermal cells contain abundant yolk reserves (Takeuchi et al., 2009), and glycerol derived from yolk lipids is likely used as a substrate for gluconeogenesis. Previous studies in other vertebrates commonly showed that the gluconeogenic tissue are closely associated with yolk reserve (Furukawa et al., 2024; Kodama et al., 2024; Shimizu et al., 2024; 2025). In contrast, epidermis of *X*. *tropicalis* is derived from the ectoderm and lacks substantial yolk, suggesting a novel role of developmental gluconeogenesis in this tissue. Interestingly, a previous study reported a concurrent upregulation of urea cycle and gluconeogenesis in 5-day-old tadpoles, raising the hypothesis that this metabolic shift contributes to metamorphosis (Konno, 2023). In this article, mRNA encoding CPSI, a key enzyme in urea cycle, was found in the gill and tail of the tadpole. Because these tissues are rich in ectoderm-derived cells, association of urea cycle and gluconeogenesis in these tissues will be interesting to address in the future. At present, however, no further evidence exists supporting the above hypothesis. Future studies should lead to a deeper understanding of physiological roles of gluconeogenesis during amphibian development, particularly in the epidermis. Collective insights into developmental gluconeogenesis in various animals may contribute to a better understanding of their functional roles and evolutionary diversity throughout vertebrate lineages.

## Materials and methods

The western clawed frog (*Xenopus tropicalis*) strains Nigerian BH and BH2 were provided by Hiroshima University Amphibian Research Center. *X. tropicalis* were kept in the laboratory of School of Marine Bioscience, Kitasato University. Two to three adult individuals were kept in each 5-L tanks containing dechlorinated tap water. They were fed with fish feed (Catfish feed; Kyorin, Hyogo, Japan) three times a week to satiation, and the water was exchanged 2 h after feeding.

### Mating

Male and female individuals were injected with 100 U pregnant mare serum gonadotropin (PMSG; ASKA Pharmaceutical, Tokyo, Japan). After 2 weeks, they were injected with 10 U human chorionic gonadotropin (hCG; ASKA Pharmaceutical), followed 24 h later by the injection of 100 U hCG. The frogs were then kept in tanks containing 0.01 x Marc’s modified Ringer’s (MMR) buffer (1 mM NaCl, 0.02 mM KCl, 0.01 mM MgSO_4_, 0.02 mM CaCl_2_, 0.05 mM HEPES, 1 μM EDTA; pH 7.8) and left in dark. Spawning began approximately 4–5 h after the final injection. The eggs were dejellied in cysteine solution (4% L-cysteine, 0.1 x MMR, 0.1 N NaOH; pH 7.8) and washed 0.1x MMR buffer three times. Unfertilized eggs (at 0 hpf), embryos (at 3, 6, 12, and 18), and larvae (at 24, 36, 48, 72, and 96 hpf) were sampled with reference to the developmental stages described in Xenbase (http://www.xenbase.org/entry/). All experimental procedures were approved by institutional animal care and use committee (approval nos. MB2024-010, MB2025-011) and genetic engineering safety committee (no. 5435) of Kitasato University.

### Metabolite analysis

Metabolites were measured using liquid chromatography-quadrupole time-of-flight mass spectrometry (LC-MS). Thirty unfertilized eggs (at 0 hpf), embryos (at 3, 6, 12, and 18), and larvae (at 24, 36, 48, 72, and 96 hpf) were collected into 1.5 ml tubes (30 individuals per tube, n=5-7), anesthetized with 0.2% MS-222, and rinsed with water. Subsequently, 200 μL of 100% methanol and 5 μL of internal standard solution (L-methionine sulfone, 2- (*N*-morpholino) ethanesulfonic acid, D-camphor-10-sulfonic acid, 5 mM) were added, and the samples were homogenized. Next, 100 μL water and 200 μL of chloroform were added. The tubes were thoroughly mixed and incubated on ice. After 10 min, samples were centrifuged at 13,000 rpm for 5 min at 4℃, and the supernatants were collected. The aqueous phase was analyzed using an LC system (Shimadzu, Kyoto, Japan) coupled to a TripleTOF 5600+ mass spectrometer (AB Sciex, Framingham, MA). Amino acid and phosphorylated sugars were measured by Shodex HILICpak VT-50 2D (Resonac, Tokyo, Japan). Other metabolites were measured by Shodex HILICpak VG-50 2D (Resonac). Detailed analytical conditions are listed in Table S1. For glycogen analysis, 10 µL of each sample was aliquoted into two tubes and dried using a centrifugal evaporator. For one tube (control group), 10 µL of 0.5% acetic acid was added and the samples were incubated at 4 °C for 2 h. For another tube (glucoamylase treatment), 10 µL of a mixed solution containing 0.5% acetic acid and glucoamylase (final concentration 300 U/mL) was added, followed by incubation at 37 °C for 2 h. These samples were further subjected to methanol/chloroform extraction and analyzed by LC-MS. Glycogen content was determined by measuring glucose levels in glucoamylase-treated and untreated samples, and the difference between them was defined as glycogen content. Data was analyzed using PeakView and MultiQuant software (AB Sciex). Metabolite concentrations were quantified using calibration curves generated from serial dilutions of standard solutions with known concentrations.

### Isotope tracing

After anesthesia, *X. tropicalis* larvae at 24 hpf were cut into three pieces. L-alanine-^13^C3 (Merck, Darmstadt, Germany), sodium L-lactate-^13^C3 (Cambridge Isotope Laboratories, Tewksbury, MA, USA), glutamate-^13^C5 (Taiyo Nippon Sanso, Tokyo, Japan), glutamine-^13^C5 (Cambridge Isotope Laboratories), glycerol-^13^C3 (Taiyo Nippon Sanso) were diluted to 5 mM with L-15 medium. Larvae were immersed for 6 h in the media containing each ^13^C metabolic substrate or in control medium without ^13^C substrate (n=6). After incubation, samples were transferred to 1.5 mL tubes and homogenized in 100 μL 100% ethanol and 2.5 μL internal standard solution. Metabolites were extracted as described in the previous section. Samples were analyzed LCMS-8045 system (Shimadzu). Phosphorylated sugars and amino acids were measured using a Shodex HILICpak VT-50 2D (Resonac). UDP-glucose and UDP-GlcNAc were measured using a Shodex HILICpak VG-50 2D (Resonac). Metabolites in the TCA cycles were measured using a Intrada Organic Acid (Imtakt Corporation, Kyoto, Japan). Detailed analytical condition is listed in Table S2. The detection mode (positive or negative), collision energy (CE), and *m/z* values for precursor and product ions of each metabolite were determined using the optimization program in LabSolutions (Shimazu). The *m/z* values of precursor/product ions and CE for each metabolite are listed in Table S3. Isotopic enrichment of metabolite was expressed as (M+n)/(M+0) *100, where M represents the unlabeled mass of each metabolite.

### Real-time quantitative PCR

The total RNA was extracted from the samples at unfertilized eggs (at 0 hpf), embryos (at 3, 6, 12, and 18), and larvae (at 24, 36, 48, 72, and 96 hpf) (N=7) with TRI reagent (Molecular Research Center, Cincinnati, OH). The RNA treated with DNase Ⅰ (Roche Diagnostics, Basel, Switzerland) and 3 μg of total RNA was reverse transcribed with ReverTraAce qPCR RT Kit (Toyobo, Osaka, Japan). qPCR was performed with Thermal Cycler Dice Real-time SystemⅡ (Takara Bio, Shiga, Japan) and Luna Universal qPCR Master Mix (New England Biolabs, Ipswich, MA). Targeted genes related to gluconeogenesis and glycogen metabolism were: glucose-6-phosphatase (G6PC), *g6pc1*/*g6pc1.2*/*g6pc1.3*; glycerol-3-phosphate dehydrogenase (GPD), *gpd1*/*gpd1l*/*gpd2*; fructose-1,6-phosphatase (FBP), *fbp1*; Lactate dehydrogenase (LDH), *ldha*/*ldhb*; phosphoenolpyruvate carboxykinase (PEPCK), *pck1*/*pck2*; glycogen synthase (GYS), *gys1*/*gys2*; glycogen phosphorylase (PYG), *pygb*/*pygl*/*pygm*. The primers used were listed in Table S4. After the PCR amplification, the specificity of the gene was confirmed by dissociation curve analysis. For quantification, plasmids containing the target gene fragments were used as standard solutions at known concentrations (10^1^∼10^7^ copies/μL), and standard curves were drawn by parallel amplifications of these standards. The expression levels in the samples were calculated based on the standard curves.

### RNA probe synthesis

For *in situ* hybridization analysis, DIG-labeled RNA probes were synthesized. First, targeted cDNA fragments were amplified with the primers listed in Table S4 and ligated into the pGEM T-Easy vector (Promega, Madison, WI). The cDNA fragments were amplified by PCR with M13 primers and purified with a Fast Gene Gel/PCR Extraction Kit (Nippon, Genetics, Tokyo, Japan). The purified cDNA fragments were transcribed *in vitro* with T7 or SP6 RNA polymerases (Roche, Diagnostics, Mannheim, Germany) in the presence of digoxigenin (DIG) labeling Mix (Roche Diagnostics). The resulting RNA probes were digested with DNaseⅠand purified with the RNeasy MinElute cleanup Kit (Qiagen, Hilden, Germany).

### *In situ* hybridization

The larvae of 24, 48, 96 hpf were fixed in 4% paraformaldehyde (PFA) in phosphate-buffered saline (PBS) prepared with diethylpyrocarbonate (DEPC)-treated water (PBS-DEPC) overnight at 4℃. Fixed larvae were embedded in paraffin and sliced into 5 μm. After deparaffinization, the samples were digested with 2 μg/mL proteinase K in PBS-DEPC for 30 min at room temperature and postfixed in 4% PFA in PBS-DEPC for 15 min. The samples were acetylated twice for 10 m with 0.1 M triethanolamine (pH 8.0) containing 0.25% acetic anhydride (Hayashi et al., 1978). After prehybridizing the samples in hybridization mixture {HM^+^; 50% formaldehyde, 5x saline-sodium citrate buffer (SSC), 0.01% Tween 20, 50 mg/ml heparin and 0.5 mg/ml yeast tRNA; Thisse and Thisse, 2008} at 70 ℃ for 2 h, the samples were hybridized with the RNA probes at the concentration of 65 ng / 200 μl in HM^+^ overnight at 70℃. For *pck1*, prehybridization and hybridization were carried out at 75℃. Next day, the samples were washed in 0.2x SSC and PBS. The sample were blocked in 5% skim milk in PBS for 1h at 4℃ and incubated for 1 h at 4℃ in blocking solutions with anti-DIG antibody (Roche Diagnostics) diluted 1:10,000 mRNA signals were visualized by BCIP/NBT Alkaline Phosphatase Substrate Kit (Vector Laboratories). Coloration signals were observed under a microscope (BZ-X 700; Keyence, Osaka, Japan), and digital images were captured.

### Immunofluorescence staining for glycogen

The larvae of 72 hpf were fixed in 4% PFA in PBS-DEPC overnight at 4℃. Fixed larvae were embedded in paraffin and sliced into 5 μm. After deparaffinization, the sample were treated with a mixture of 0.5% acetic acid and glucoamylase (0.5% acetic acid; 300 U/ml glucoamylase), while control samples were treated with acetic acid only and incubated at 37℃ for 2 h. After wash in water and PBS, The sample were blocked in PBS containing 1x Western Blocking Reagent (Roche Diagnostics) for 1 h at 4℃. Then, samples were incubated with anti-glycogen antibody (Baba, 1993.) diluted 1: 1000 with PBS containing 1x Western Blocking Reagent overnight at 4℃. Next day, the samples were washed in water and PBS, and incubated in Alexa Fluor 568-conjucated goat anti-mouse IgM (ab175702; Abcam, Cambridge, UK) diluted in 1: 2000 for 1 h at 4℃. The samples were washed with water and stained with DAPI (Dojindo Laboratories, Kumamoto, Japan). Samples were observed by LSM800 (Carl Zeiss Microscopy, Mannheim, Germany). The excitation / detection wavelengths were: Alexa Flour 563 (568 nm/579-603 nm), DAPI (405 nm/400-505 nm).

### Generation of *fbp1* G0 KO embryos using CRISPR/Cas9

Four scrambled gRNAs lacking a target sequence and four guide RNAs (gRNAs) targeting *X. tropicalis fbp1* were generated by *in vitro* transcription (Wu et al., 2018). Sequences of the scaffold, scrambled, and *fbp1*-specific primers used for gRNA synthesis are listed in Table S4. Male were induced for spermiation using the same protocol as for mating. Excised testes were suspended in L-15 medium containing 10% fetal bovine serum (FBS) and homogenized to prepare a sperm suspension. Unfertilized eggs were collected from hormone-induced females and fertilized by adding the sperm suspension. Cas9-gRNA ribonucleoprotein (RNP) complexes were prepared as described previously (Nakayama et al., 2014) by mixing Cas9-NLS protein (2 μg) with gRNA mix (400 ng) at a 1:1 ratio and incubating at 37℃ for 10 min. After fertilization, the jelly coat was removed, and 1 nL of Cas9 RNP complex was microinjected into embryos at the one- to two-cell stage.

### Analysis of genome editing efficiency

Genomic DNA was extracted from *fbp1* G0 KO and SC at 24 hpf using the alkaline lysis method (Truett et al., 2000). The target region was amplified by PCR, followed by a second-round PCR using primers containing unique barcode sequences. Sequencing libraries were prepared using the barcoded PCR products and the Ligation Sequencing Kit (SQK-LSK114; Oxford Nanopore Technologies, oxford, UK). Primers are listed in Table S4. Sequencing was performed on a MinION Mk1B device with a Flongle Flow Cell (FLO-FLG114; Oxford Nanopore Technologies, Oxford, UK). Raw signal data were basecalled using Dorado (v7.9.8) integrated within the MinKNOW software (v25.05.14), employing the Super-accurate (SUP) model. Read quality scores were obtained from the sequencing summary file generated during basecalling, and only high-quality reads with a Q score >20 were retained for downstream analysis by extracting the corresponding reads from the FASTQ files. The mutation frequency (Indel efficiency) was quantified using Geneious Prime (2025.1.3). Individual reads were mapped to the wild-type reference sequence using the minimap2 algorithm. Insertions or deletions (Indels) of 1 bp or more located within a 10 bp window surrounding the predicted sgRNA cleavage site (3 bp upstream of the PAM; Jinek et al., 2012) were counted. The mutation rate was calculated as the ratio of the total number of reads containing indels to the total number of valid mapped reads. To eliminate potential sequencing errors, single-nucleotide substitutions and minor variants outside the target window were excluded from the analysis.

### Histological analysis of brain tissue by HE staining

At 48 hpf, the *fbp1* G0 KO and SC larvae were fixed in 4% PFA, embedded in paraffin, and cross sections of 5 μm-thickness were prepared. The sections were subjected to hematoxylin and eosin (HE) staining. Sections at comparable anatomical levels containing the mesencephalon were selected for the analysis. For both *fbp1* G0 KO and SC groups (n=6 each), mesencephalon area and ventricle area were measured using Fiji. The ratio of ventricle area to mesencephalon area was then calculated.

### Statistics

All numerical data are presented as mean ± standard error (SE). Differences among groups were considered statistically significant at *p*< 0.05. Metabolite abundance and qPCR data were analyzed by one-way analysis of variance (ANOVA) followed by Tukey’s multiple comparison test. The results of metabolite tracing experiments were analyzed by one-way ANOVA followed by Dunnett’s multiple comparison test. For the metabolite tracing experiments and ventricle/mesencephalon area ratios in G0 KO larvae, comparisons between two groups were performed using Welch’s *t*-test. A *p*-value < 0.05 was considered statistically significant. All statistical analyses were performed using GraphPad Prism 11 (GraphPad Software, Boston, MA).

## Acknowledgment

The *X. tropicalis* strain “Nigerian BH/BH2” was kindly provided by Hiroshima University Amphibian Research Center (RRID: SCR_019015) through the National BioResource Project (NBRP) of the MEXT, Japan.

---

**Table S1. Conditions of LC-MS**

**Table S2. Conditions of LC-MS**

**Table S3. LC-MS parameters used for data acquisition**

**Table S4. Primers used in this study**

**Fig. S1. Histological analysis during development by HE staining.**

Embryos and larvae of *X. tropicalis* at 12, 18, 24, 48, 72 and 96 hpf were subjected to hematoxylin and eosin (HE) staining to examine tissue at different developmental stages. A) 12 hpf; B) 18 hpf; C) 24 hpf; D, E) 48 hpf; F, G) 72 hpf; H, I) 96 hpf. Scale bars represent 200 μm in A-D, F, and H, and 100 μm in E, G and I. ar, archenteron; br, brain; cg, cement gland; dg, digestive gland; ey, eye; ec, ectoderm; en, endoderm; ep, epidermis; ga, ganglion; gi, gill; he, heart; in, intestine; li, liver; me, mesoderm; mg, midgut; mo, mouth; mu, muscle; nc, notochord; no, nose; nt, neural tube; pd, pronephric; sm, somite.

**Fig. S2. LC-MS based isotope tracing reveals gluconeogenesis activities in *X. tropicalis* larvae.**

The horizontal axes represent ^13^C-labeled substrates (alanine, lactate, glutamine, glutamate, glycerol) and the vertical axes represent the ratio of mass (M) +3 to M+0 isotopologues of the detected metabolites. Data are presented as mean ±SE (N=5-6). Different letters indicate statistically significant differences among substrates (*p*<0.05), as determined by one-way analysis of variance (ANOVA) followed by Dunnett’s multiple comparison test. DHAP, Dihydroxyacetone Phosphate; S7P, Sedoheptulose-7-phosphate; Glucose-6-P, Glucose-6-phosphate; Glucose-1-P, Glucose-1-phosphate; Fructose-6-P, Fructose-6-phosphate; Glycerol-3-P, Glycerol-3-phosphate; GAP, Glyceraldehyde-3-phosphate; PEP, Phosphoenolpyruvate.

**Fig. S3. LC-MS based isotope tracing reveals TCA cycles activities in *X. tropicalis* larvae.**

(A) Schematic overview of the TCA cycles illustrating the incorporation of ^13^C-labeled substrates. (B) The horizontal axes represent ^13^C-labeled substrate (alanine, lactate, glutamine, glutamate, glycerol) and the vertical axes represent the ratio of mass (M) +2 or 3, 4, 5 to M+0 isotopologues of the detected metabolites. Data are presented as mean ±SE (N=5-6). Different letters indicate statistically significant differences among substrates (*p*<0.05), as determined by one-way analysis of variance (ANOVA) followed by Dunnett’s multiple comparison test.

**Fig. S4. *In situ* hybridization for gluconeogenic and glycogen metabolism genes at 24 hpf.**

For all transcripts, sense probes (right) were used as negative control for antisense probes (left), which showed true signals. Positive signals are indicated by black arrowheads and tissue labels. br, brain; en, endoderm; ep, epidermis; he, heart; mu, muscle. Scale bars represent 250 μm (*in situ* hybridization, gluconeogenic genes), 500 μm (*in situ* hybridization, glycogen metabolism genes, i.e., *gys1*, *gys2*, *pygb*, *pygm*, *pygl*), and 200 μm (HE staining).

**Fig. S5. *In situ* hybridization for gluconeogenic and glycogen metabolism genes at 48 hpf.**

For all transcripts, sense probes (right) were used as negative control for antisense probes (left), which showed true signals. Positive signals are indicated by black arrowhead and tissue labels. br, brain; en, endoderm; ep, epidermis; he, heart; mu, muscle. Scale bars represent 250 μm (*in situ* hybridization, gluconeogenic genes), 500 μm (*in situ* hybridization, glycogen metabolism genes, i.e., *gys1*, *gys2*, *pygb*, *pygm*, *pygl*), and 200 μm (HE staining).

**Fig. S6. *In situ* hybridization for gluconeogenic and glycogen metabolism genes of 96 hpf.**

For all transcripts, sense probes (right) were used as negative control for antisense probes (left), which showed true signals. Positive signals are indicated by black arrowheads. ep, epidermis; in, intestine; he, heart; li, liver; mu, muscle. Scale bars represent 250 μm (*in situ* hybridization, gluconeogenic genes), 500 μm (*in situ* hybridization, glycogen metabolism genes, i.e., *gys1*, *gys2*, *pygb*, *pygm*, *pygl*), and 200 μm (HE staining).

**Fig. S7. Whole-mount *in situ* hybridization for gluconeogenic genes at 24 hpf.**

For all transcripts, sense probes (right) were used as negative control for antisense probes (left), which showed true signals. Positive signals are indicated by black arrowheads and tissue labels. ep, epidermis; sm, somite. Scale bars represent 300 μm.

**Fig. S8. Whole-mount *in situ* hybridization for gluconeogenic genes at 48 hpf.**

For all transcripts, sense probes (right) were used as negative control for antisense probes (left), which showed true signals. Positive signals are indicated black arrowheads and tissue labels. br, brain; ep, epidermis; mu, muscle. Scale bars represent 1 mm.

**Fig. S9. Whole-mount *in situ* hybridization for gluconeogenic genes at 96 hpf.**

For all transcripts, sense probes (right) were used as negative control for antisense probes (left), which showed true signals. Positive signals are indicated black arrowheads and tissue labels. br, brain; ep, epidermis. Scale bars represent 1 mm.

**Fig. S10. Immunofluorescence staining for glycogen at 72 hpf.**

Glycogen localization was confirmed at 72 hpf. The bottom panels show the glucoamylase (GA) treated (+) negative control. White arrowheads in non-GA treated (-) panels indicate the position of glycogen (red), which is not found from GA (+) counterparts. Blue signals (DAPI) indicate nuclei. Scale bars indicated 200 μm (A), 100 μm (B). Tissues are labeled as follows: in, intestine; li, liver; mu, muscle.

**Fig. S11. Allelic composition of SC and G0 KO larvae based on amplicon analysis by nanopore sequencing.**

Allelic composition in representative G0 KO larvae. The pie chart displays the proportions of on-target indels and WT alleles. In addition, the three most frequent alleles are shown individually, with indel size and frequency (%) indicated for each allele. Deletions are indicated by the Δ, while combined deletion/insertion events are represented as, for example, △ 4/ + 9. When multiple alleles exhibited the same frequency as the third-ranked allele, all such alleles are additionally shown.

**Fig. S12. Top allelic composition of individual G0 KO larvae based on amplicon analysis by nanopore sequencing.**

The top five most frequent alleles in each larva are shown individually. For each allele, indel size is indicated. Sequences are aligned to the WT sequence, with deletions indicated by hyphens (–) and insertions or substitutions shown in color.

**Fig. S13. Metabolomic and isotope tracer analyses using LC–MS in SC and *fbp1* G0 KO larvae.**

(A, B) Metabolites involved in gluconeogenesis/glycolysis, glycogen metabolism, the pentose phosphate pathway, and the hexosamine biosynthetic pathway were analyzed. (A) metabolite content (nmol/individual). (B) Ratios of mass (M) +*n* (*n* = 2, 3, 4, 5) to M+0 isotopologues for each metabolite. Each dot represents an individual replicate (5 larvae), and bars indicate mean ±SE (n = 6). Statistical comparisons between two groups were performed using Welch’s *t*-test (\**p* < 0.05, \*\**p* < 0.01, \*\*\**p* < 0.001, \*\*\**p* < 0.0001). G6P, glucose-6-phosphate; G1P, glucose-1-phosphate; F6P, fructose-6-phosphate; UDP-GlcNAc, UDP-N-acetylglucosamine; S7P, sedoheptulose-7-phosphate; Asp, aspartate; Gln, glutamine.

**Fig. S14. Histological analysis of brain tissue by hematoxylin and eosin (HE) staining in SC and G0 KO larvae.**

Representative HE-stained images of SC (top) and G0 KO (bottom) larvae (n=6). Scale bars represent 200 µm.

## References

Agius, L. (2015). Role of glycogen phosphorylase in liver glycogen metabolism. Mol. Aspects. Med. 46: 34–45.

Assheton, R. (1896). Notes on the ciliation of the ectoderm of the amphibian embryo. J. Cell. Sci. s2-38 (152): 465–484.

Baba, O. (1993). [Production of monoclonal antibody that recognizes glycogen and its application for immunohistochemistry] [in Japanese]. Kokubyo Gakkai Zasshi 60: 264–287.

Bansil, R., Turner, B. S. (2018). The biology of mucus: Composition, synthesis and organization. Adv. Drug. Deliv. Rev. 124: 3–15.

Bennett, E. P., Mandel, U., Clausen, H., Gerken, T. A., Fritz, T. A., Tabak, L. A. (2011). Control of mucin-type O-glycosylation: a classification of the polypeptide GalNAc-transferase gene family. J. Glycobiol. 22 (6): 736–756.

Billett, F. S., Gould, R. P. (1971). Fine structural changes in the differentiating epidermis of *Xenopus laevis* embryos. J. Anat. 108 (3): 465–480.

Bulusu, V., Prior, N., Snaebjornsson, M. T., Kuehne, A., Sonnen, K. F., Kresse ., Stein, F., Schultz, C., Sauer, U., Aulehla, A. (2017). Spatiotemporal Analysis of a Glycolytic Activity Gradient Linked to Mouse Embryo Mesoderm Development. Dev.Cell. 40 (4): 331–341.

Cao, L., Zhang, D., Chen, J., Qin, Y. Y., Sheng, R., Feng, X., Chen, Z., Ding, Y., Li. M., Qin, Z. H. (2017). G6PD plays a neuroprotective role in brain ischemia through promoting pentose phosphate pathway. Free. Radic. Biol. Med. 112: 433–444.

Chaveroux, C., Sarcinelli, C., Barbet, V., Belfeki, S., Barthelaix, A., Ferraro-Peyret, C., Lebecpue, S., Renno, T., Bruhat, A., Fafournoux, P., Manié, N. S. (2016). Nutrient shortage triggers the hexosamine biosynthetic pathway via the GCN2-ATF4 signalling pathway. Sci. Rep. 6: 27278.

Czajewski, I., van Aalten, D. M. F. (2023). The role of O-GlcNAcylation in development. Dev. 150 (6): dev.201370.

Deberardinis, R. J., Mancuso, A., Daikhin, E., Nissim, I., Yudkoff, M., Wehrli, S., Thompson, C. B. (2007). Beyond aerobic glycolysis: Transformed cells can engage in glutamine metabolism that exceeds the requirement for protein and nucleotide synthesis. Proc. Natl. Acad. Sci. U.S.A. 104 (49):19345–19350.

Deberardinis, R. J., Chandel, N. S. (2016). Fundamentals of cancer metabolism. Sci. Adv. 2 (5).

Denzel, M. S., Antebi, A. (2015). Hexosamine pathway and (ER) protein quality control. Curr. Opin. Cell. Biol. 33: 14–18.

Dubaissi, E., Rousseau, K., Lea, R., Soto, X., Nardeosingh, S., Schweickert, A., Amaya, E., Thornton, D. J., Papalopulu, N., (2014). A secretory cell type develops alongside multiciliated cells, ionocytes and goblet cells, and provides a protective, anti-infective function in the frog embryonic mucociliary epidermis. Development 141 (7): 1514–25.

Dubaissi, E., Rousseau, K., Hughes, G, W., Ridley, C., Grencis, R. K., Roberts, I. S., Thornton, D. J. (2018). Functional characterization of the mucus barrier on the *Xenopus tropicalis* skin surface. Proc. Natl. Acad. Sci. U.S.A. 115 (4): 726–731.

Dworkin, M. B., Dworkin-Rastl, E. (1990). Regulation of carbon flux from amino acids into sugar phosphates in *Xenopus* embryos. Dev. Biol. 138 (1): 177–187.

Figlak, K., Williams, G., Bertolini, M., Paus, R., Philpott, M. P. (2021). Human hair follicles operate an internal Cori cycle and modulate their growth via glycogen phosphorylase. Sci. Rep. 11: 20761.

Fogotto, F., Maxfield, F. R. (1994). Changes in yolk platelet pH during Xenopus laevis development correlate with yolk utilization. A quantitative confocal microscopy study. J. Cell. Sci. 107 (12): 3325–3337.

Furukawa, F., Aoyagi, A., Sano, K., Sameshima, K., Goto, M., Tseng, Y. C., Ikeda, D., Lin, C. C., Uchida, K., Okumura, S., Yasumoto, K., Jimbo, M., Hwang, P. P. (2024). Gluconeogenesis in the extraembryonic yolk syncytial layer of the zebrafish embryo. PNAS. Nexus. 3 (4): 125.

Goncalves J., Wan, Y., Guo, X., Rha, K., LeBoeuf, B., Zhang, L., Estler, K., Garcia, L. R. (2020). Succinate dehydrogenase regulated phosphoenolpyruvate carboxykinase sustains copulation fitness in aging *C. elegans* males. iScience. 23 (4): 100990.

Grainger, R. M. (2014). *Xenopus tropicalis* as a model organism for genetics and genomics: past, present and future. Mol. Biol. 917: 3–15.

Hakim, A. M., Moss, G., Scuderi, D. (1980). The pentose phosphate pathway in brain during development. Biol Neonate. 37 (1-2): 15–21.

Harland, R. M., Grainger, R. M. (2011). Xenopus Research: Metamorphosed by Genetics and Genomics. Trends. Genet. 27 (12): 507–515.

Harris, J. M., Esain, V., Frechette, G. M., Harris, L. J., Cox, A. G., Corte, M., Garmaas, K., Carroll, K. J, Cutting, C. C., Khan, T., Elks, P. M., Renshaw, S. A., Dickinson, B. C., Chang, C. J., Murphy, M. P., Paw, B. H., Heiden, M. G. V., Goessling, W., North, T. E. (2013). Glucose metabolism impacts the spatiotemporal onset and magnitude of HSC induction in vivo. Blood. 121 (13): 2483–2493.

Hayashi, S., Gillam, I. C., Delaney, A. D., Tener, G. M. (1978). Acetylation of chromosome squashes of Drosophila melanogaster decreases the background in autoradiographs from hybridization with [125I]-labeled RNA. J. Histochem. Cytochem. 26 (8): 677–679.

Hellsten, U., Harland, R. M., Gilchrist, M. J., Hendrix, D., Jurka, J., Kapitonov, V., Ovcharenko, I., Putnam, N. H., Shu, S., Taher, L., Blitz, I. L., Blumberg, B., Dichmann, D. S., Dubchak, I., Amaya, E., Detter, J. C., Fletcher, R., Gerhard, D. S., Goodstein, D., Graves, T., Grigoriev, I. V., Grimwood, J., Kawashima, T., Lindquist, E., Lucas, S. M., Mead, P. E., Mitros, T., Ogino, H., Ohta, Y., Poliakov, A. V., Pollet, N., Robert, J., Salamov, A., Sater, A. K., Schmutz, J., Terry, A., Vize, P. D., Warren, W. C., Wells, D., Wills, A., Wilson, R. K., Zimmerman, L. B., Zorn, A. M., Grainger, R., Grammer, T., Khokha, M. K., Richardson, P. M., Rokhsar, D. S. (2010). The Genome of the western clawed frog Xenopus tropicalis. Sci. 328 (5978): 633–636.

Hui, S., Ghergurovich, J. M., Morsher, R. J., Jang, C., Teng, X., Lu, W., Esparza, L. A., Reva, T., Zhan, L., Guo, J. Y., White, E., Rabinowitz, J. D. (2017). Glucose feeds the TCA cycle via circulating lactate. Nature 551:115–118.

Holeček M. (2023) Roles of malate and aspartate in gluconeogenesis in various physiological and pathological states. Metab. 145.

Jensen, P. J., Gitlin, J. D., Carayannopulos, M. O. (2006). GLUTI deficiency links nutrient availability and apoptosis during embryonic development. Biol. Chem. 281 (19): 13382–13387.

Jinek, M., Chylinski, K., Fonfara, I., Hauer, M., Doudna, J. A., Charpntier, E. (2012). A programmable dual-RNA-guided DNA endonuclease in adaptive bacterial immunity. Sci. 337 (6097): 816–821.

Jorgensen, P., Steen, J. A. J., Steen, H., Kirschner, M. W. (2009). The mechanism and pattern of yolk consumption provide insight into embryonic nutrition in *Xenopus*. Dev. 136 (9): 1539–1548.

Kimmel, C. B., Law, R. D. (1985). Cell lineage of zebrafish blastomeres. II. Formation of the yolk syncytial layer. Dev. Biol. 108 (1): 86–93.

Kodama, T., Watanabe, S., Kayanuma, I., Sasaki, A., Kurokawa, D., Baba, O., Jimbo, M., Furukawa, F., (2024). Gluconeogenesis during development of the grass puffer (*Takifugu niphobles*). Comp. Biochem. Physiol. A Mol. Integr. Physiol. 295: 111663.

Konno, N. (2023). Simultaneous activation of genes encoding urea cycle enzymes and gluconeogenetic enzymes coincides with a corticosterone surge period before metamorphosis in *Xenopus laevis*. Dev. Growth Differ. 65 (1): 6–15.

Lechenault, H., Mellinger, J. (1993). Dual origin of yolk nuclei in the lesser spotted dogfish, *Scyliorhinus canicula* (Chondrichthyes). J. Exp. Zool. 265 (6): 669–678.

Liu, H., Wang, S., Wang, J., Guo, X., Song, Y., Fu, K., Gao, Z., Liu D., He, W., Yang, L. L. (2025). Energy metabolism in health and disease. Signal. Transduct. Target. Ther. 10: 69.

Nakayama, T., Blitz, I. L., Fish, M. B., Odeleye, A. O., Manohar, S., Cho, K. W. Y., Grainger, R. M. (2014). Cas9-based genome editing in *Xenopus tropicalis*. Methods. Enzymol. 546: 355–375.

Needham, J. (1927). The carbohydrate metabolism in amphibian embryogenesis. Q. J. Exp. Physiol. 18 (2): 153–154.

Niewkoop, P. D., and Faber, J. (1994). Normal Table of Xenopus laevis (Doudin): a systematical and chronological survey of the development from the fertilized egg till the end of metamorphosis. Garland publishing.

Marr, L., Biswas, D., Daly, L. A., Browning, C., Vial, S. C. M., Maskell, D. P., Hudson, C., Bertrand, J. A., Pollard, J., Ranson, N. A., Khatter, H., Eyers, C. E., Sakamoto, K., Zeqiraj, E. (2022). Mechanism of glycogen synthase inactivation and interaction with glycogenin. Nat. Commun. 13: 3372.

Oginuma, M., Moncuquet, P., Xiong, F., Karoly, E., Chai, J., Guevorkian, K., Pourquié, O. (2017). A gradient of glycolytic activity coordinates FGF and Wnt signaling during elongation of the body axis in amniote embryos. Dev. Cell. 40 (4): 342–353.

Pantaleon, M., Scott, J., Kaye, P. L. (2008). Nutrient sensing by the early mouse embryo: hexosamine biosynthesis and glucose signaling during preimplantation development. Biol. Reprod. 78 (4): 595–600.

Patel, J. H., Ong, D. J., Williams, C. R., Callies, L. K., Wills, A. E. (2022). Elevated pentose phosphate pathway flux supports appendage regeneration. Cell. Rep. 41 (4): 111552.

Patra, K. C., Hay, N. (2014). The pentose phosphate pathway and cancer. Trends. Biochem. Sci. 39 (8): 347–54.

Quigley, I. K., Stubbs, J. L., Kintner, C. (2011). Specification of ion transport cells in the *Xenopus* larval skin. Dev. 138 (4): 705–714.

Rizzo, A. M., Gornati. R., Galli. C., Bernardini, G., Berra, B., (1994). Cholesterol, triacyclglycerols and phospholipids during *Xenopus* embryo development. Cell. Biol. Int. 18 (11): 1085–1090.

Roach, P. J., Depaoli-Roach, A. A., Hurley, T. D., Tagliabracci, V. S. (2012). Glycogen and its metabolism: some new developments and old themes. Biochem. J. 441 (3): 763–787.

Ruan, D., Hu, T., Yang, X., Mo, X., Ju, Q. (2025). Lactate in skin homeostasis: metabolism, skin barrier, and immunomodulation. Front. Immunol. 16: 1510559.

Sahoo, B., Srivastava, M., Katiyar, A., Ecelbarger, C., Tiwari, S., (2023). Liver or kidney: Who has the oar in the gluconeogenesis boat and when? World. J. Diabetes. 14 (7): 1049–1056.

Shah, M. A., Xie, X., Marek, R., Stundl, J., Ingo, B., Šindelka, R., Rzepkowska, M., Saito, T., Pšenička, M. (2024). Sturgeon gut development: a unique yolk utilization strategy among vertebrates. Front. Cell. Dev. Biol. 12: 1358702.

Shimizu, M., Takagi, W., Sakai, Y., Kayanuma, I., Furukawa, F. (2024). Gluconeogenesis in the yolk syncytial layer-like tissue of cloudy catshark (*Scyliorhinus torazame*). Physiol. Rep. 12 (11).

Shimizu, M., Matsuzaki, T., Kobayashi, K., Aoyagi, A., Kaneko, M., Nagasawa, T., Hiraoka, K., Furukawa, F., (2025). Gluconeogenesis and glycogen metabolism during development in sterlet *Acipenser ruthenus* and bester. Fish. Sci. 91: 881–890.

Showell, C., Conlon, F. L. (2013). Natural mating and tadpole husbandry in the western clawed frog *Xenopus tropicalis*. Cold. Spring. Harb. Protoc. 2009 (9).

Stincone, A., Prigione, A., Cramer, T., Wamelink, M. M. C., Campbell, K., Cheung, E., Olin-Sandoval, V., Grüning, N. M., Krüger, A., Alam, M. T., Keller, M. A., Breitenbach, M., Brindle, K. M., Rabinowitz, J. D., Ralser, M. (2015). The return of metabolism: biochemistry and physiology of the pentose phosphate pathway. Biol. Rev. 90 (3): 927–63.

Takeuchi, M., Takahashi, M., Okabe, M., Aizawa, S., (2009). Germ layer patterning in bichir and lamprey; an insight into its evolution in vertebrates. Dev. Biol. 332 (1): 90–102.

Territo, P. R., Smits, A. W., (1998). Whole-body composition of *Xenopus laevis* larvae: implications for lean body mass during development. J. Exp. Biol. 201 (Pt7): 1013–1022.

Trinkaus, J. P. (1993). The yolk syncytial layer of Fundulus: its origin and history and its significance for early embryogenesis. J. Exp. Zool. 265 (3): 258–284.

Truett, G. E., Heeger, P., Mynatt, R. L., Truett, A. A., Walker, J. A., Warman, M. L. (2000). Preparation of PCR-quality mouse genomic DNA with hot sodium hydroxide and tris (HotSHOT). Biotechniques. 29 (1): 52–54.

Thisse, C., Thisse, B., (2008). High-resolution in situ hybridization to whole-mount zebrafish embryos. Nat. Protoc. 3: 59–69.

Vander Heiden, M. G., Cantley, L. C., Thompson, C. B. (2009). Understanding the warburg effect: the metabolic requirements of cell proliferation. Sci. 324 (5930): 1029–1033.

Vastag, L., Jorgensen, P., Peshkin, L., Wei, R., Rabinowitz, D. J., Kirschner, W. M., (2011). Remodeling of the metabolome during early frog development. PLoS. ONE. 6 (2): e16881.

Wrisez, F., Lechenault, H., Leray, C., Haye, B., Mellinger, J. (1993). Fate of yolk lipid in an oviparous elasmobranch fish, *Scyliorhinus canicula* (L.). Fish. Physiol. Biochem. 315–322.

Wu, R. S., Lam, I. I., Clay, Hilary., Duong, D. N., Deo, R. C., Coughlin, S. R. (2018). A rapid method for directed gene knockout for screening in G0 zebrafish. Dev. Cell. 46 (1): 112–125.

Zahn, N., James-Zorn, C., Ponferrada, V. G., Adams, D. S., Grzymkowski, J., Buchholz, D. R., Nascone-Yoder, N. M., Horb, M., Moody, S. A., Vize, P. D., Zorn, A. M. (2022). Normal table of *Xenopus* development: a new graphical resource. Dev. 149 (14): dev200356.

