## Supplementary figures and images for "Gluconeogenesis and glycogen metabolism in the epidermis and endoderm of *Xenopus tropicalis* embryos and larvae"

### Supplementary Fig. S1

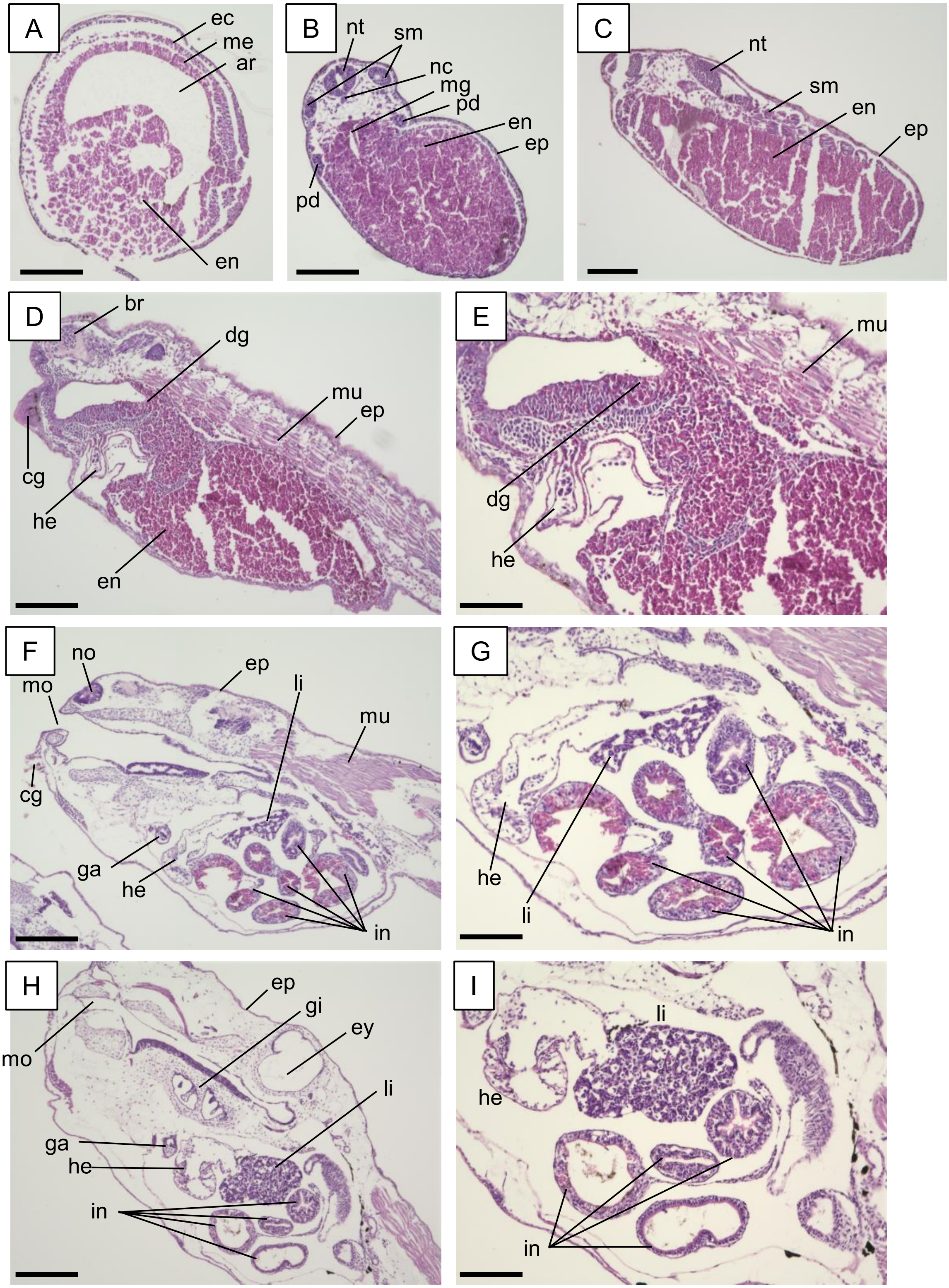

### Supplementary Fig. S2

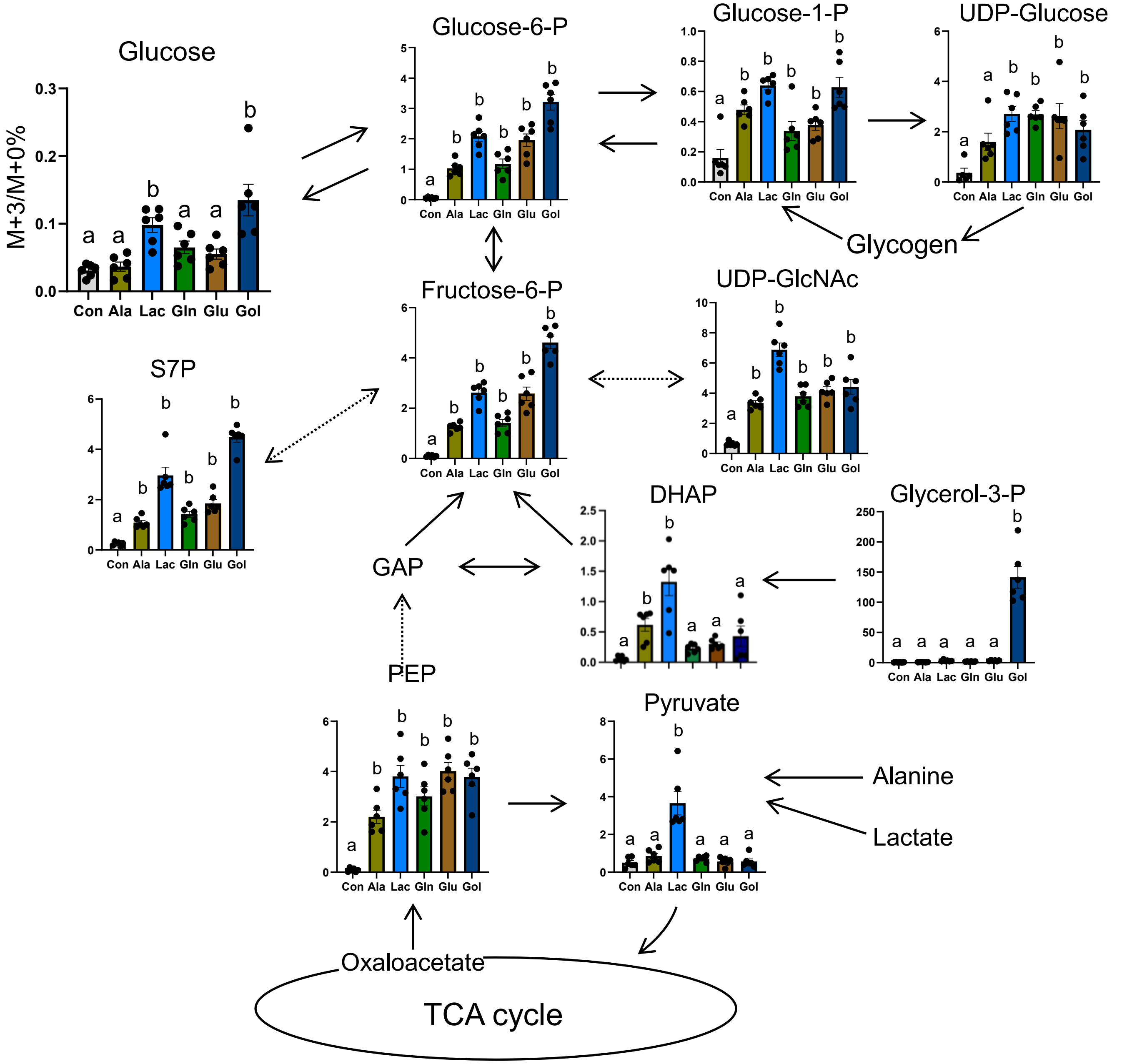

### Supplementary Fig. S3

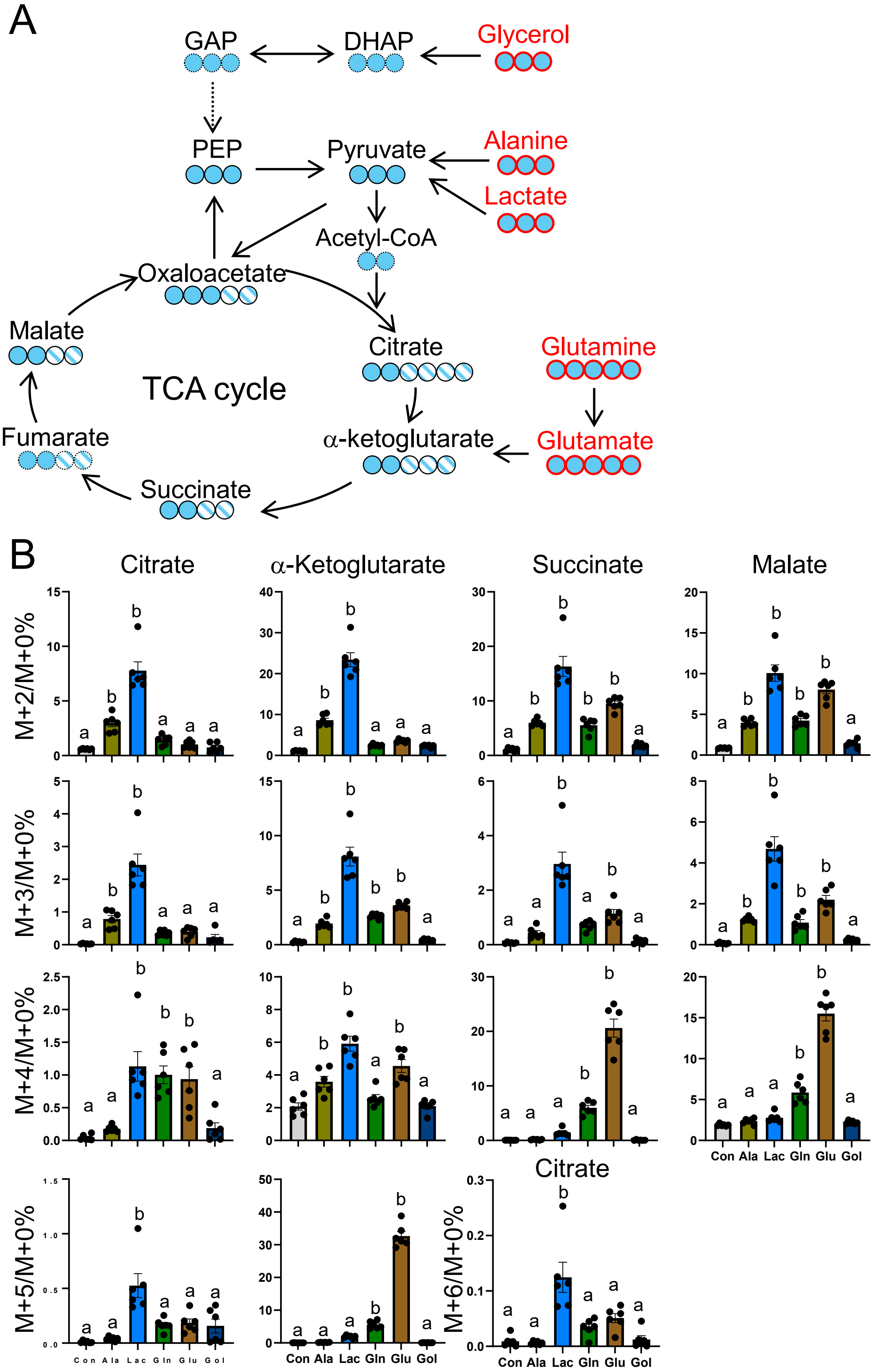

### Supplementary Fig. S4

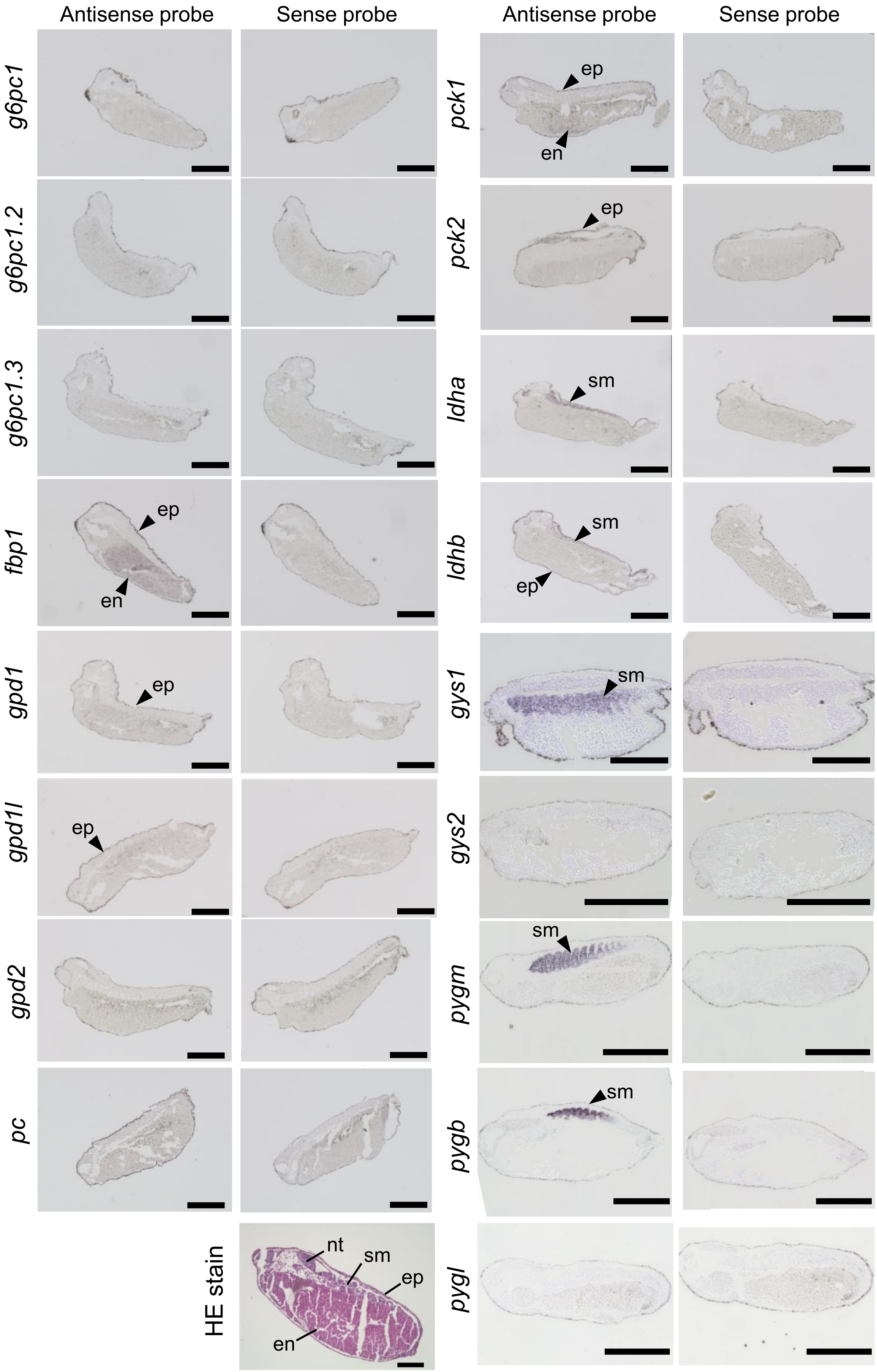

### Supplementary Fig. S5

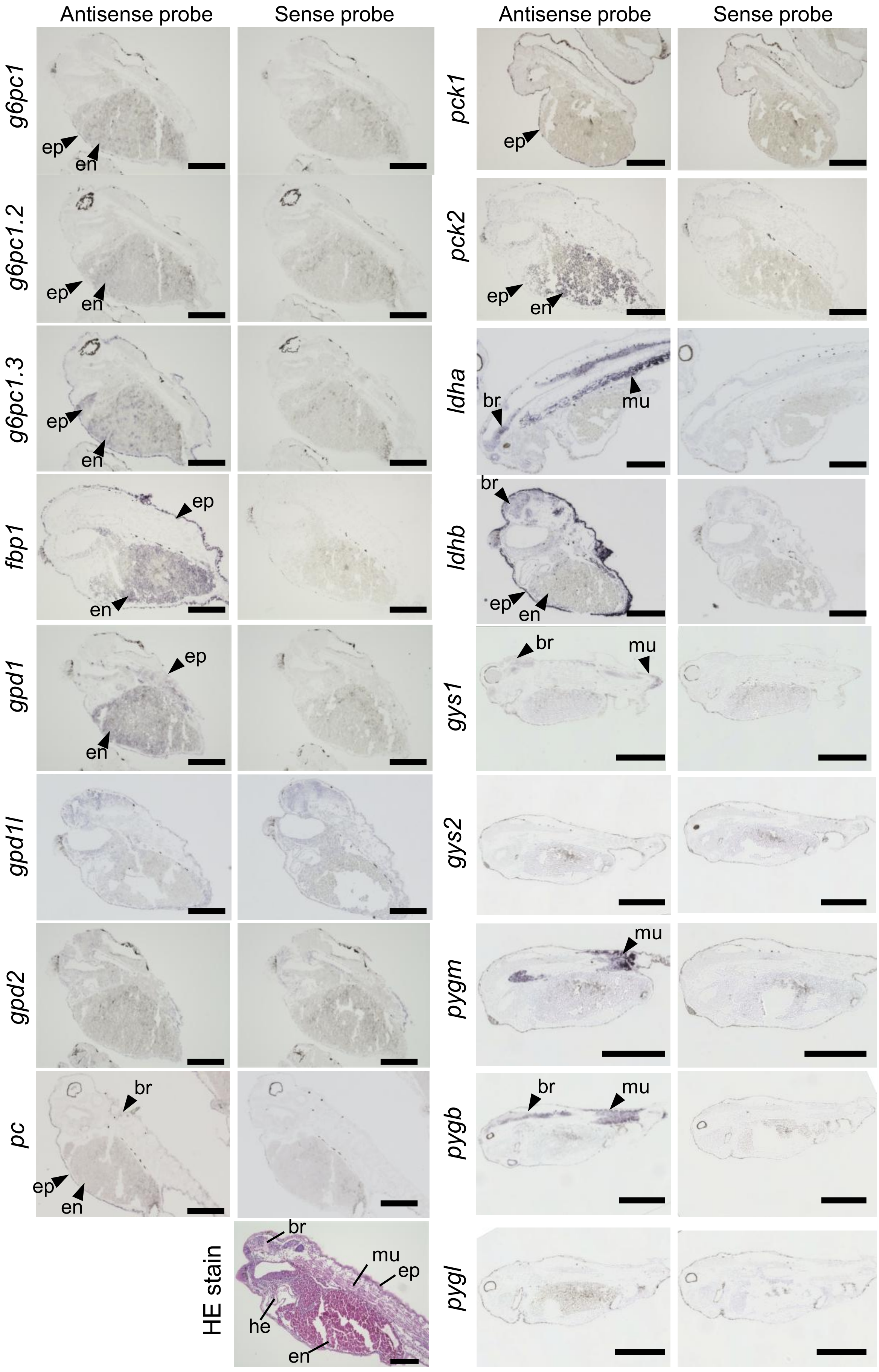

### Supplementary Fig. S6

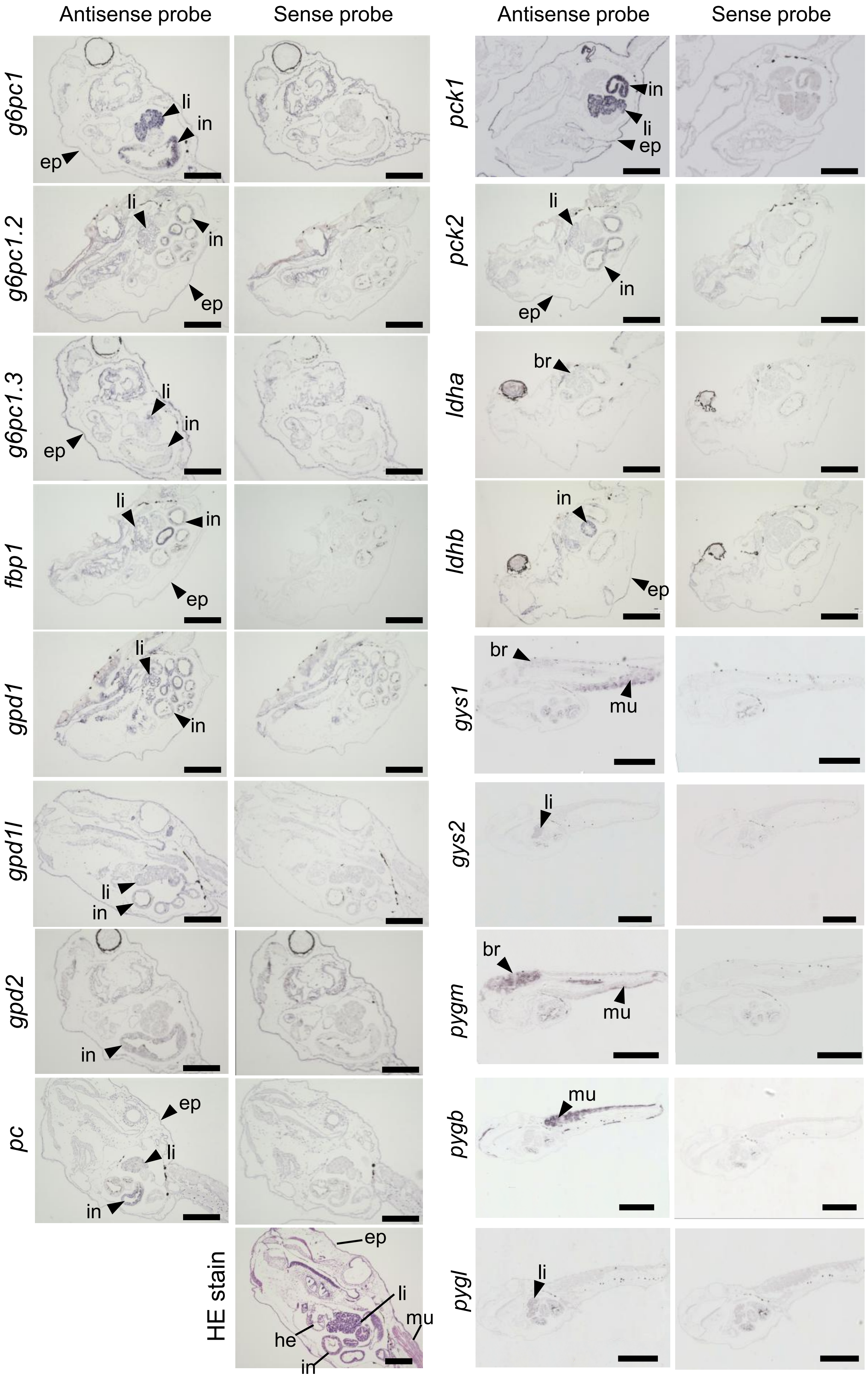

### Supplementary Fig. S7

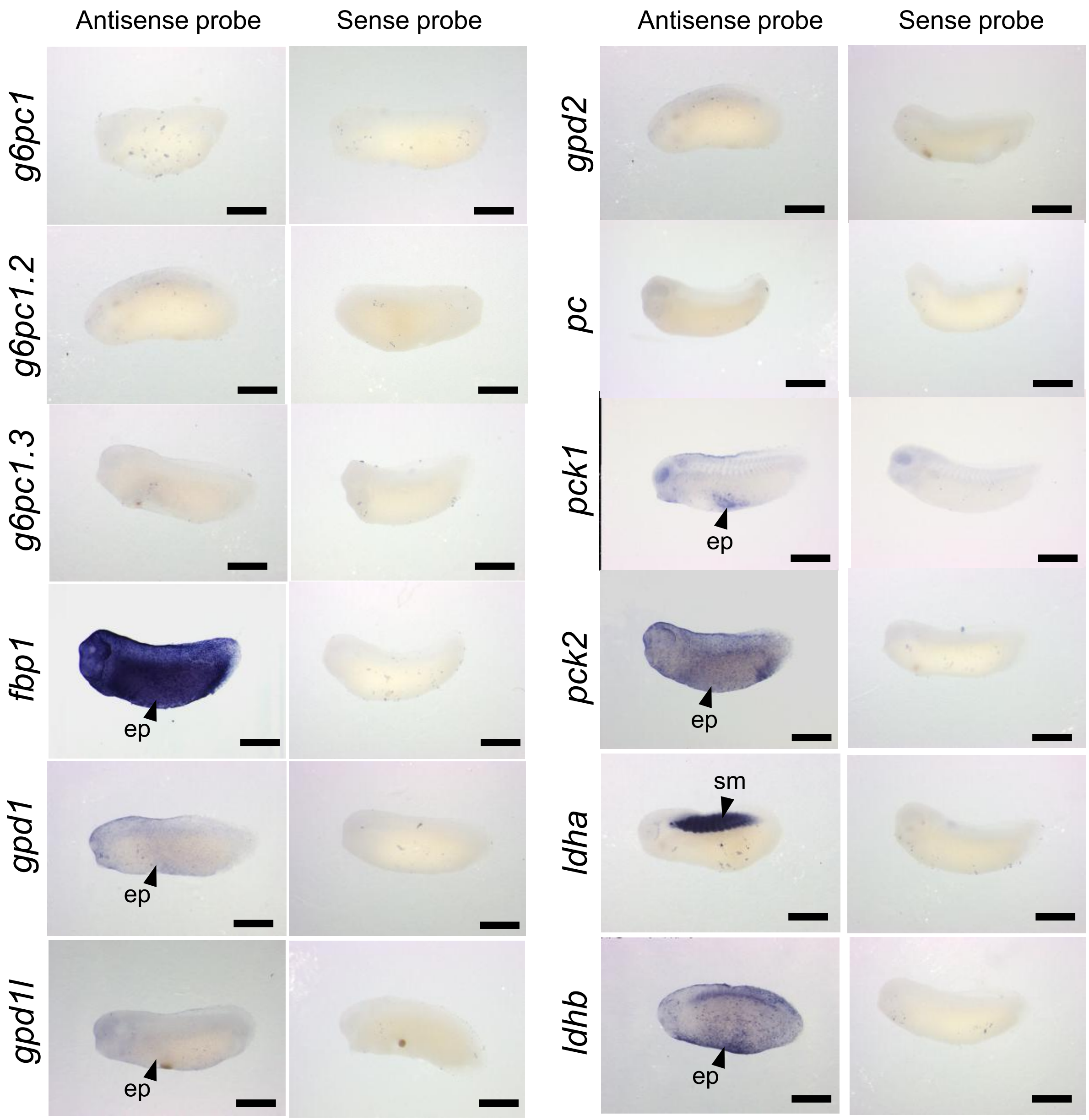

### Supplementary Fig. S8

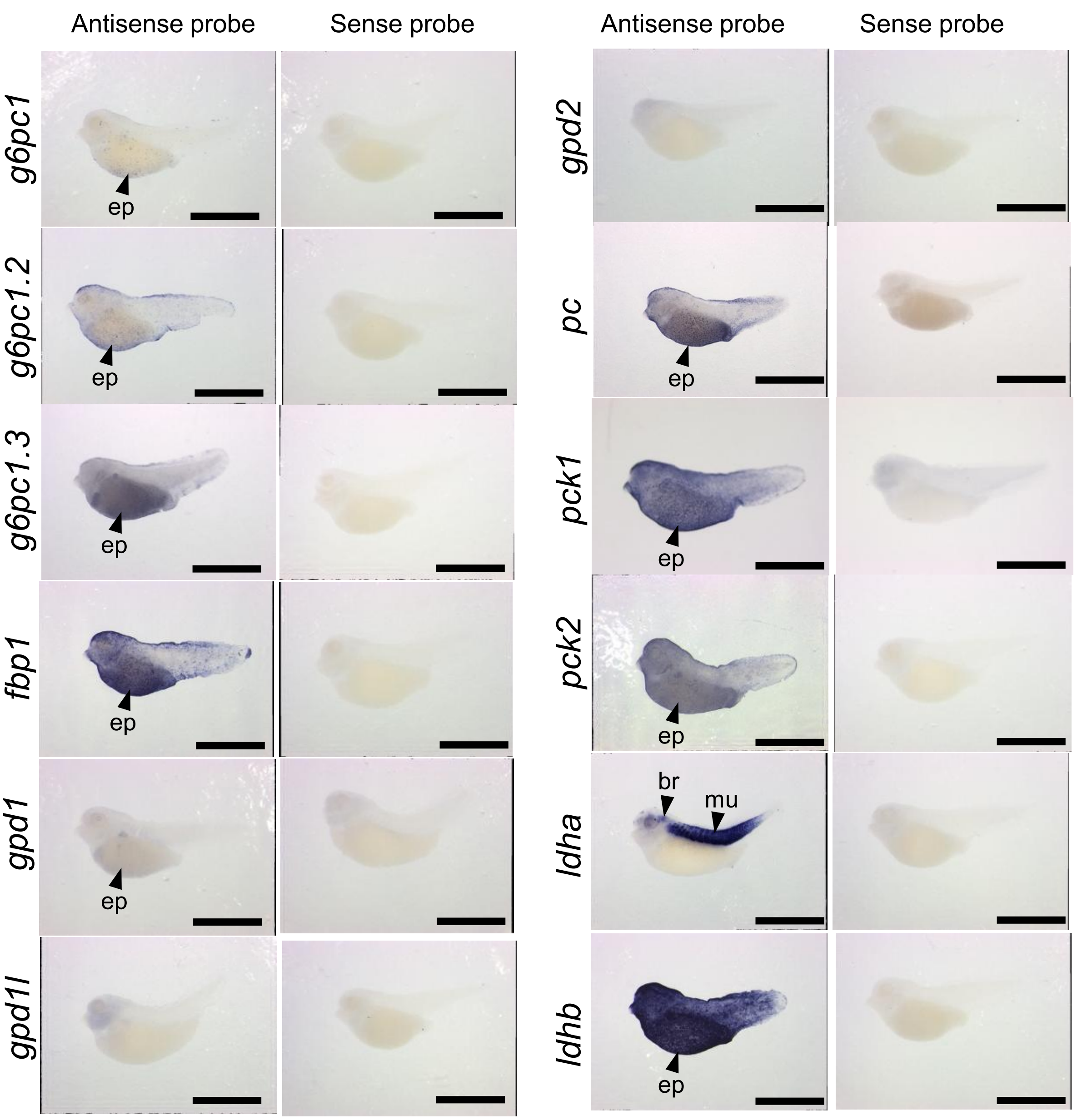

### Supplementary Fig. S9

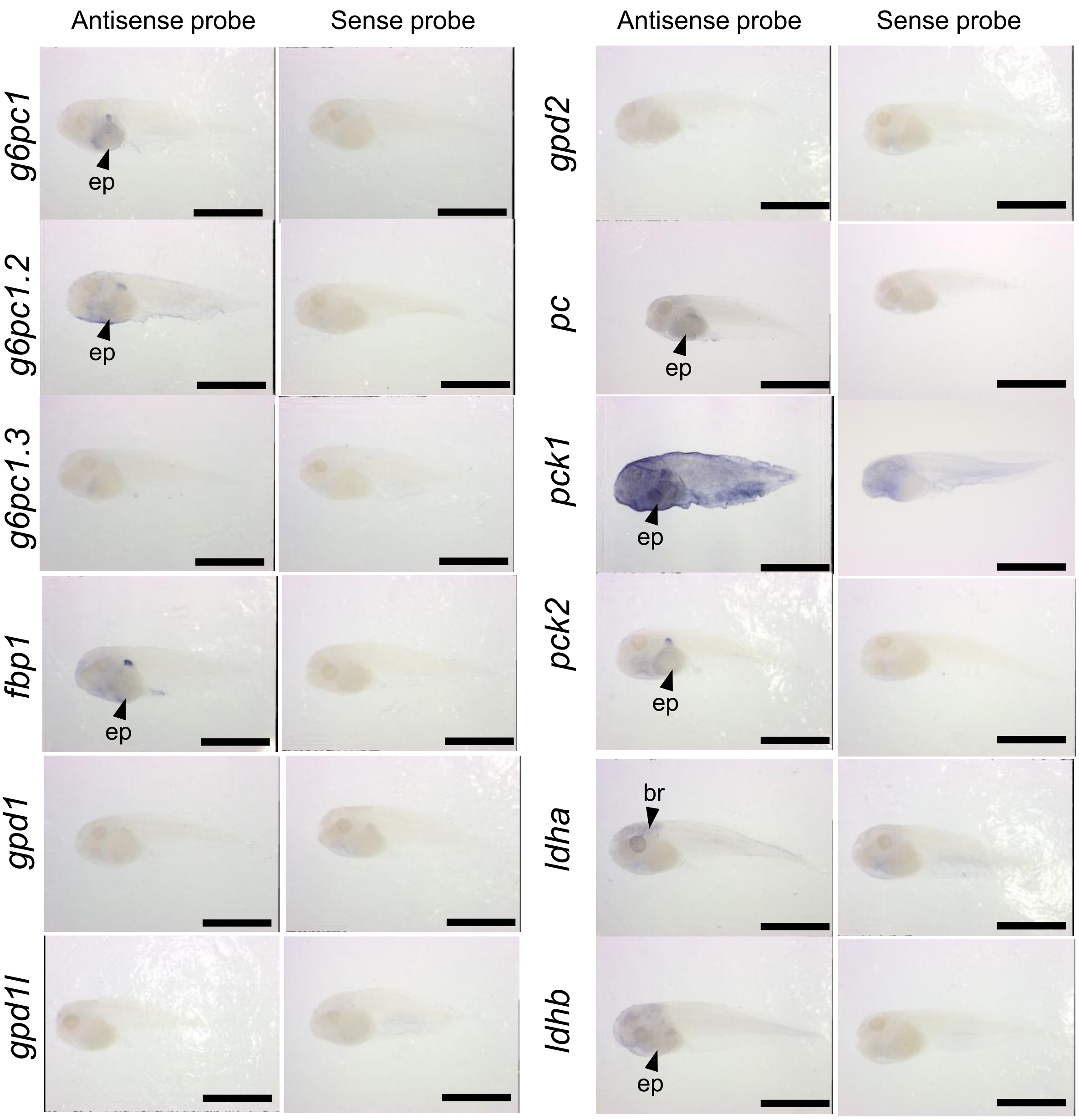

### Supplementary Fig. S10

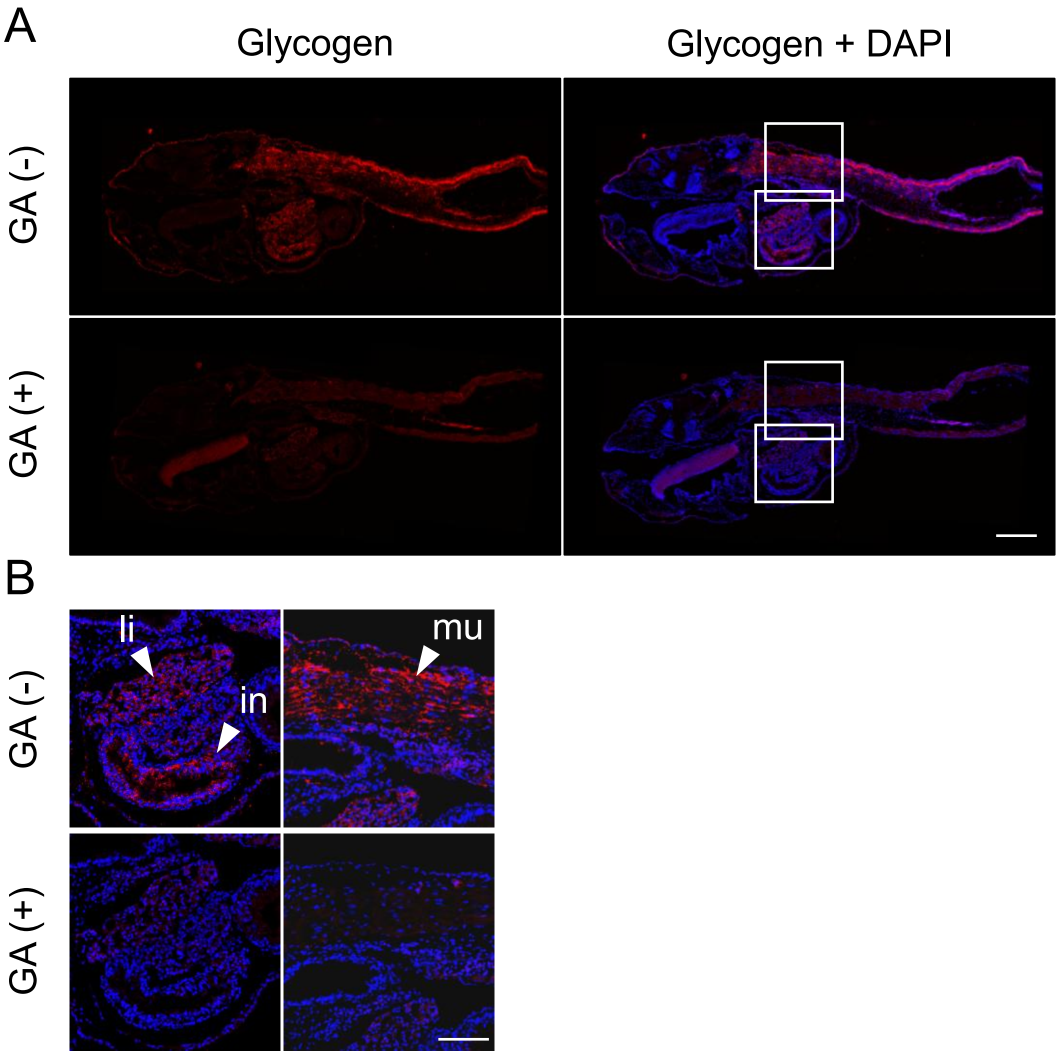

### Supplementary Fig. S11

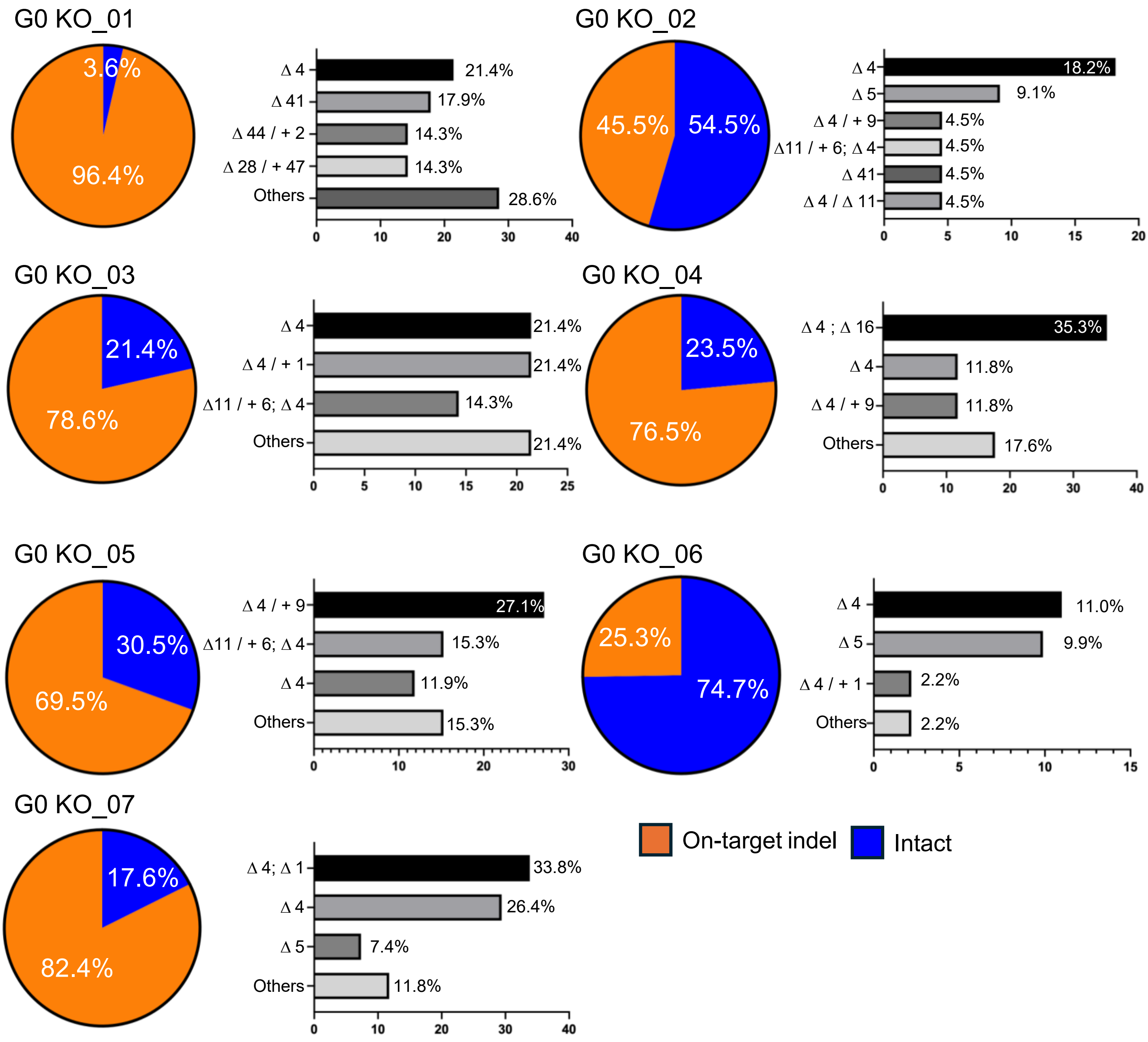

### Supplementary Fig. S12

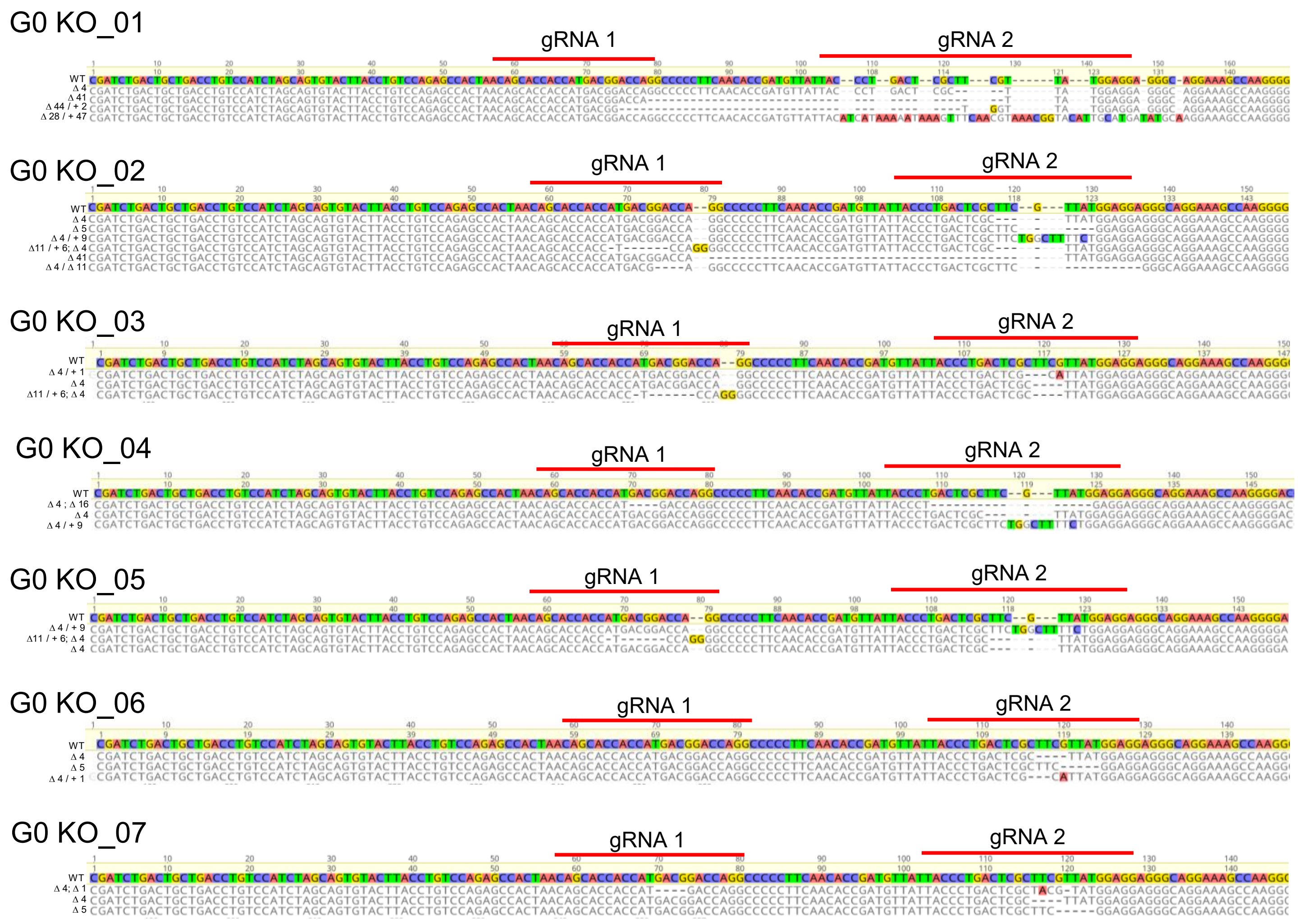

### Supplementary Fig. S13

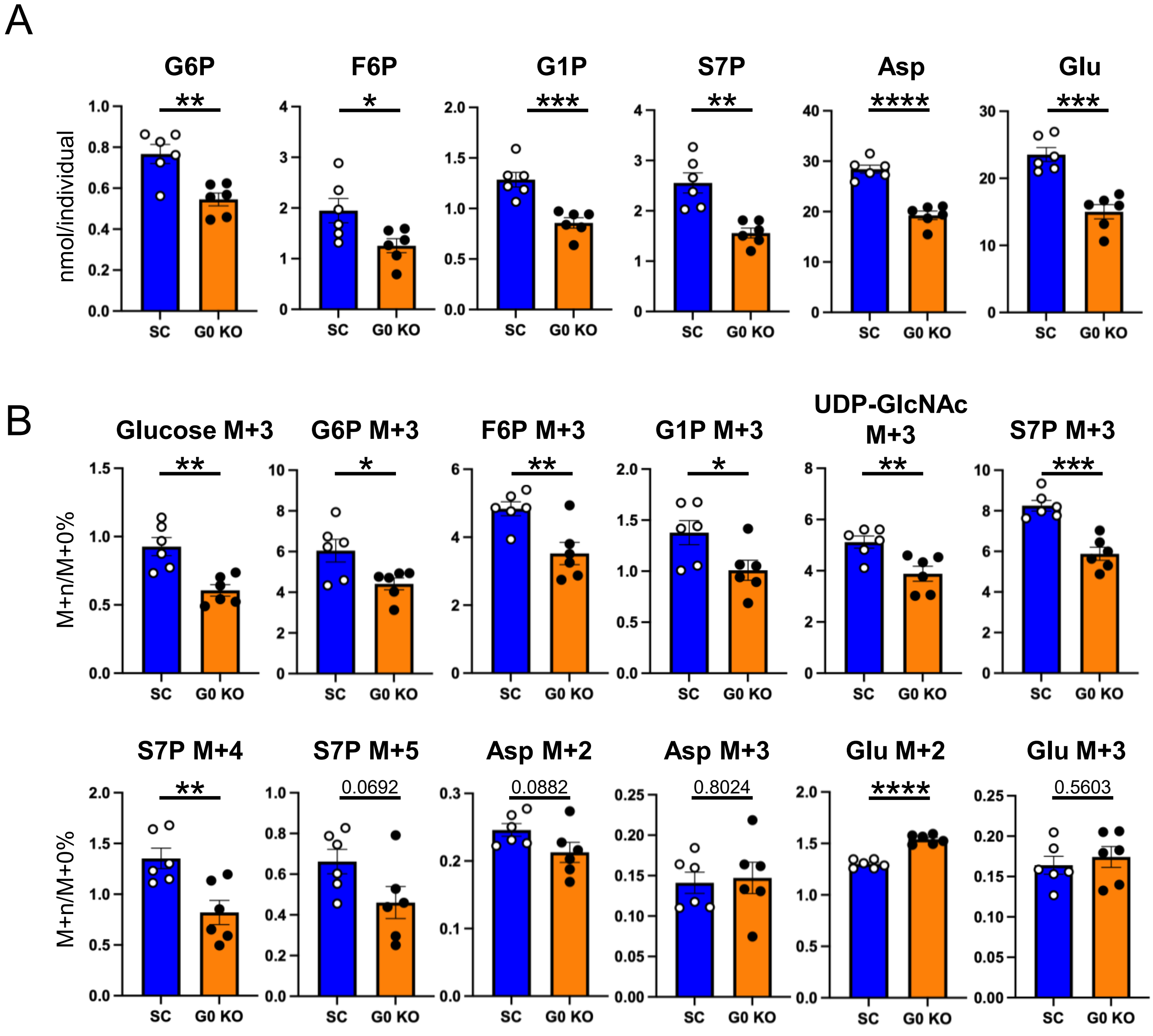

### Supplementary Fig. S14

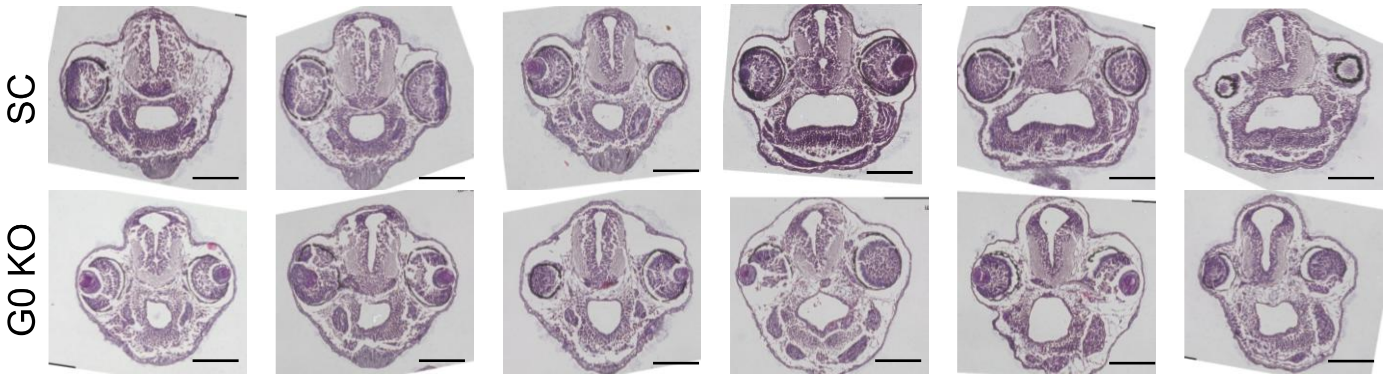
